# Disc-Toroid Hybrid Lipid Nanoparticles for Efficient Drug Encapsulation and Subcutaneous Delivery

**DOI:** 10.1101/2025.07.20.665764

**Authors:** Zanelle van Niekerk, Rima Nuwayhid, Stefaniya Gaydarova, Eva Bittrich, Natalia Makarova, Susanne Boye, Petr Formanek, Christo Tzachev, Jan C. Simon, Sandra Franz, Albena Lederer

## Abstract

The development of an effective for subcutaneous or intradermal injection drug delivery requires systems with improved bioavailability and biocompatibility. Systematic physicochemical and biological interrogation of carnauba-wax/red-palm-oil lipid nanoparticles (LNPs) stabilised with d-α-tocopheryl-PEG-1000-succinate and polysorbate-40 shows that purposeful matrix engineering yields a robust sub-50 nm carrier for under-skin delivery. Cryo-TEM and SAXS uncover a disc-toroid hybrid morphology dominated by 30–40 nm particles with toroidal/disc shape – an advantageous biconcave geometry to enhance surface-to-volume ratio and is expected to accelerate enzymatic erosion after injection. Orthogonal analytics (AF4-MD, DLS, MALS, WAXS) confirmed that loading with quinine or dihydroartemisinin leaves size and crystallinity unchanged while delivering encapsulation efficiencies of approximately 90 % and long-term particle stability up to 18 months at 4 °C. Red-palm oil and the dual-surfactant corona act synergistically to suppress bimodality and narrow size distribution compared with single-component controls. Short-term viability assays in keratinocytes, fibroblasts and macrophages showed no cytotoxicity even at ≥1 % (w/v) lipid, underscoring excellent biocompatibility. Fluorescein-labelled LNPs injected into ex vivo human skin traversed the dermis and hypodermis, while only nanomolar lipid concentrations appeared in the receiver medium, indicating a sustained local depot. Collectively, these insights link composition, structure and performance, positioning wax-based disc-toroid LNPs as a flexible platform for high-load delivery of small-molecule or biopharmaceutical therapeutics via minimally invasive under-skin administration.

## INTRODUCTION

Lipid-based nanoparticles (LNPs) have evolved from first-generation liposomes into a versatile delivery platform that now supports mRNA vaccines, RNA-interference medicines, long-acting small-molecule depots, and intracellular imaging agents.^1–3^ The LNP family extends beyond liposomes to include nanostructured liquid and solid particles. Solid lipid nanoparticles have matured from a conceptual replacement for oil-in-water emulsions into a versatile nanoplatform that combines the regulatory familiarity of lipids with the colloidal control required for modern drug delivery.^4^ Unlocking their full potential, however, demands a deeper, quantitative understanding of how composition, shape, size distribution, and internal architecture govern nano-bio interactions and therapeutic performance.^5, 6^ Typical solid LNPs are 50-300 nm dispersions produced by high-pressure homogenisation or melt emulsification and stabilised by non-ionic or polymeric surfactants.^7^ Their fully or partially crystalline cores confer long-term physical stability, protect chemically labile drugs, and enable manufacturing without organic solvents.^8^ Yet the same crystallinity reduces lattice defects, making high drug load and controlled release difficult, but the loss of payload during cooling or storage remains the dominant limitation.^9^

A widely adopted strategy to overcome this limitation is to blend a high-melting solid lipid with a low-melting liquid lipid or functional oil. Carnauba wax, a complex mixture of long-chain esters that melts at 81-86 °C and is already used in food and topical products, has proven particularly attractive.^10^ When combined 1:1 with liquid esters or triglycerides, the melting enthalpy of the matrix drops from approximately 160 to 70 J g⁻¹, indicating a less-ordered lattice that can accommodate more guest molecules without compromising colloidal stability.^10^ Such binary matrices routinely reach entrapment efficiency > 90 % for hydrophobic actives and remain physically stable for almost 30 days, even under accelerated conditions.^11^

Particle morphology adds another, often overlooked level of control. In practice, wax–oil LNPs have been reported as anisotropic-shaped particles by small-angle X-ray scattering (SAXS), features that correlate with rapid cellular internalisation and depot-like behaviour in skin.^12^ Shape engineering, therefore, offers a complementary handle to composition and size distribution.

Against this backdrop, the particles chosen for this study are LNPs with a purpose-built binary matrix composed of carnauba wax (solid) and red-palm oil (liquid) stabilised by a mixed surfactant shell of d-α-tocopheryl PEG 1000 succinate (TPGS) and polysorbate 40 with the commercial name Cellinject ™.^13^ The phase-inversion protocol yields nanoparticles with hydrodynamic diameters of 35-55 nm, sizes already within the parenteral window for subcutaneous or intradermal injection. The carnauba wax furnishes mechanical integrity, while the unsaturated triglycerides of red-palm oil introduce lattice imperfections analogous to classic nanostructured lipid carriers, maximising free volume for guest molecules.

These LNPs show two distinct properties apart from conventional solid LNPs. First, the particles display *zero* detectable release of encapsulated fluorophore in aqueous media; payload liberation is initiated exclusively after cellular uptake and enzymatic erosion of the matrix. Second, short-term cytotoxicity assays in HaCaT keratinocytes show no loss in viability even above 1% dispersions, underlining the biocompatibility of the lipid-surfactant combination. Confocal imaging in HeLa cells further confirms rapid internalisation and nucleolar accumulation of the released dye within 60 min, with fluorescent signal persisting for > 13 h.^13^

The high homogenisation pressure used for these LNPs mirrors parameter optimisation studies on carnauba-based systems, where 900 bar and ≤ 8 cycles routinely afford particles < 260 nm with PDI < 0.18.^10^ For these particles, however, the additional fluidisation provided by red-palm oil drives the mean diameter into the sub-50 nm regime without broadening the distribution, an essential prerequisite for reproducible injectability through fine-gauge needles. In classic solid LNPs, drug expulsion accompanies lipid recrystallisation.^9^ The disrupted lattice of the LNPs investigated in this study resembles nanostructured lipid carriers and supports high association efficiencies (> 99 %) even after four weeks at 4 °C, as shown previously for rosmarinic-acid-loaded wax particles.^11^ The absence of premature release in phosphate-buffered saline (PBS) or culture medium underscores the tight molecular fit between guest and matrix. In addition, carnauba wax contributes not only a high melting point but also oxidative robustness, an advantage for unsaturated payloads.^10^

Subcutaneous injections demand carriers that resist aggregation at high lipid concentrations, traverse the extracellular matrix without burst release, and degrade into biocompatible components. A narrow size distribution and sizes below 50 nm minimise syringe force and needle clogging, while the wax core slows erosion to create a local depot analogous to long-acting polymer implants but without acidic degradation products. In skin models, carnauba-based carriers form an occlusive lipid film that limits transepidermal water loss and enhances retention in the upper dermis,^10^ suggesting that injectable depots could similarly localise the drug near the injection site.

We harness orthogonal analytics - advanced asymmetrical flow field-flow fractionation with multiple detectors (AF4-MD), dynamic (DLS) and static light scattering (MALS), and fluorescence detector as well as SAXS and cryogenic transmission electron microscopy (Cryo-TEM) - to dissect how each compositional element (solid-liquid lipid ratio, surfactant identity, payload level) shapes particle size, polydispersity, morphology and stability. We then correlate these physicochemical fingerprints with *in vitro* cytocompatibility and *ex vivo* skin penetration to delineate design rules for under-skin therapeutics. The resulting insights will inform not only the formulation of the LNPs, but also the broader development of wax-based LNPs as minimally invasive, high-loading depots for biopharmaceuticals.

## RESULTS AND DISCUSSION

### Impact of composition on size and stability

**Figure 1** summarises how the specific LNP compositions influence colloidal stability, polydispersity, and particle size. Batch dynamic light scattering (DLS) and small-angle X-ray scattering (SAXS) provided average hydrodynamic (Rh) and gyration (Rg) radii, while asymmetric flow field-flow fractionation with multi-detector (AF4-MD) read-out resolved the full-size heterogeneity.

**Figure 1:**
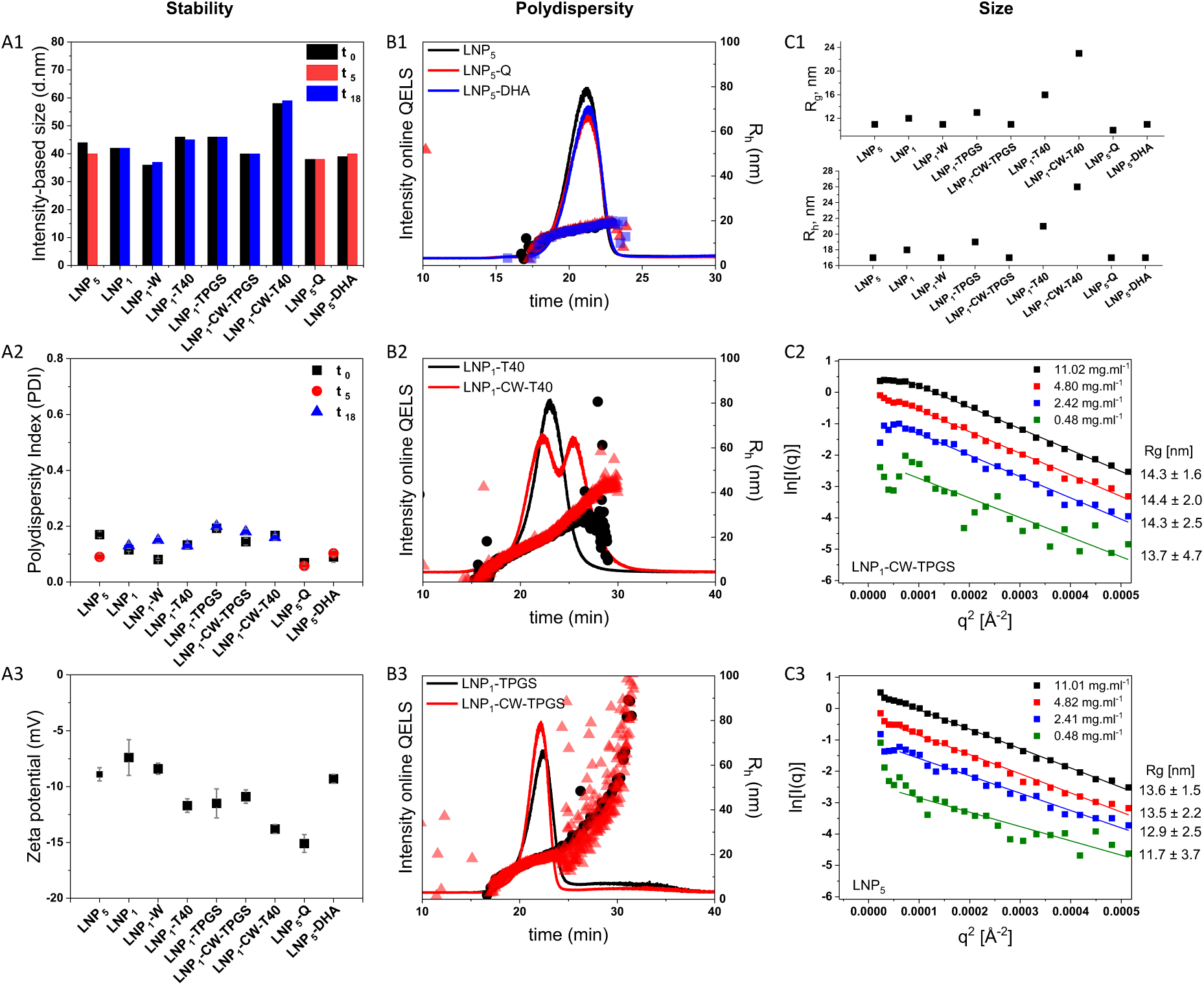
Impact of LNP composition on stability, dispersity and overall size. **(A1):** Dynamic light scattering (DLS) in batch determined intensity-based particle size (diameter) of various LNP composition formulations as a function of time over 18 months confirms stability at 4°C. **(A2):** Polydispersity index (PDI) of various LNP composition formulations as a function of time, determined using DLS. **(A3):** Zeta (ζ) potential values for the various LNP formulations. **(B1):** AF4-MD analysis of the size distribution of LNP without drug incorporation (LNP_5_) and with drug incorporation (LNP_5_-Q and LNP_5_-DHA) are compared. **(B2):** AF4-MD based comparison of two LNP formulations with the same polysorbate 40 surfactant, but different cores. LNP_1_-T40, contains a mixture of carnauba wax and red palm oil, whereas LNP_1_-CW-T40, consist only of a carnauba wax core. **(B3):** AF4-MD based comparison of two LNP formulations with the same TPGS surfactant, but different lipid cores. LNP_1_-TPGS, contains a mixture of carnauba wax and red palm oil, whereas LNP_1_-CW-TPGS, consist only of a carnauba wax core. **(C1):** AF4-MD based evaluation of the average R_h_ and R_g_ for different LNP composition formulations after separation. **(C2):** The Guinier plot from SAXS in batch evaluate the global R_g_ for four different concentrations of LNP_1_-CW-TPGS in water. **(C3):** The Guinier plot from SAXS in batch evaluate the global R_g_ for four different concentrations of LNP_5_ in water.

DLS reports Rh by converting the measured diffusion coefficient to the diameter of a hypothetical sphere with identical diffusion. In Figure 1 **A1** the intensity-weighted hydrodynamic diameters of the various formulations are plotted versus storage time of 18 months (4 °C). Most particles remain within ±10 % of their initial size, confirming good stability. Only LNP5 shows a slight decrease, consistent with surfactant desorption or gradual lipid degradation. The accompanying polydispersity indices (Figure 1 **A2**) stay below 0.20 for all samples, indicating narrow size distributions and no time-dependent broadening. LNP5’s modest fall in PDI suggests altered uniformity, whereas the small rise for LNP1-W underlines the role of the dispersion medium in particle stability. For more information, see Supplementary **Tables S1** and **S2**.

Surface charge, expressed as the zeta (ζ) potential (Figure 1 **A3; Table S3**), ranged from −8 to −15 mV. These modest negative values are sufficient to hinder aggregation through electrostatic repulsion, a conclusion supported by the absence of self-association even at 11 mg mL⁻¹ and in ultrapure water, as confirmed by SAXS (Figure 1 **C2, C3; Table S4**). Drug loading affected the ζ-potential: unloaded LNP5 measured −8.9 ± 0.6 mV, whereas quinine-loaded LNP5-Q registered −15.1 ± 0.8 mV, implying that quinine contributes additional negative charge and thereby enhances colloidal stability.

To dissect size distributions more precisely, all formulations were analysed by AF4-MD under optimised conditions (Figure 1 **B1-B3,** see also Supplementary **Table S5** and **Figure S1-S12**). Online DLS embedded in the fractograms gave Rh values between 17 and 26 nm, slightly smaller than batch DLS because larger, more highly scattering populations dominate the batch measurement. Thus, AF4 separation is essential for a complete view of particle heterogeneity, which here is derived from combined DLS, multi-angle light scattering (MALS) and UV-Vis signals.

**Figure 1 B1** compares drug-free LNP5 with its quinine-(LNP5-Q) and dihydroartemisinin-loaded (LNP5-DHA) counterparts. Each shows a single, well-defined elution peak of comparable width, yielding monomodal Rh distributions (< 17 nm ± 0.2 nm). Drug incorporation, therefore, does not alter particle size.

By contrast, lipid composition is critical. Figure 1 **B2** contrasts two polysorbate 40-stabilised systems: mixed-lipid LNP1-T40 (carnauba wax and red-palm oil) and single-lipid LNP1-CW-T40 (carnauba wax only). LNP1-CW-T40 exhibits two distinct peaks, evidencing a bimodal distribution, whereas LNP1-T40 shows a single broad peak (Rh ranged from 14 nm to over 40 nm across the eluting peak). Rh values above 20 nm across both peaks of LNP1-CW-T40 highlight heterogeneity. These data indicate that red-palm oil helps suppress polydispersity and fosters uniform particle formation.

Surfactant choice is equally decisive. Figure 1 **B3** presents TPGS-stabilised analogues: LNP1-TPGS (mixed lipids) and LNP1-CW-TPGS (carnauba wax only). Both yield single peaks with pronounced tailing toward > 80 nm and broad Rh ranging from 14 nm to over 100 nm across the elution peak. Hence, the dual-surfactant combination (Polysorbate 40 and TPGS, as in LNP5) delivers well-defined size and distribution.

Overall, incorporating both lipids and both surfactants minimises polydispersity and narrows Rh. When lipid composition is fixed, TPGS-containing particles (LNP1-TPGS) elute as a single mode, yet with higher Rh values than the Polysorbate 40 formulation (LNP1-T40).

AF4-MALS provided Rg values for particles large enough (> 10–15 nm) to scatter anisotropically (Figure 1 **C1**). These averages agree with the Rg extracted from SAXS. Because SAXS excels at very small sizes, a dilution series of LNP1-CW-TPGS and LNP5 in water (and, for completeness, in 0.9 % NaCl) was measured. No aggregation was detected, so only water data were evaluated (Figure 1 **C2, C3; Figure S21**). Guinier analysis across four concentrations gave Rg = 14 ± 2 nm for LNP1-CW-TPGS and 13 ± 2 nm for LNP5, matching the AF4-MALS results in Figure 1 **C1** and confirming that both systems share essentially the same gyration radius.

### Shape and internal composition

Cryogenic transmission electron microscopy (cryo-TEM) captures the particles in their fully hydrated state, eliminating artefacts from drying or staining. The micrographs for LNP5 (Figure 2 **A1**) and LNP1-CW-TPGS (Figure 2 **A2**) reveal two dominant morphologies. Most particles appear spherical, 30–40 nm in diameter (Figure 2 **A1.1**), yet a fraction exhibits toroidal contrast, hinting at compositional variations within the core. A minority of more elongated, high-contrast objects (thickness ≈ 7 - 9 nm; Figure 2 **A1.2**) are also evident. More micrographs of the LNPs are available in the Supplementary Information (see **Figures S13-S18**). Three-dimensional rendering (see **Figure S19**) confirms an overall disc-like shape irrespective of composition. To probe topology, crystallinity, and shape in greater detail, AF4 and X-ray scattering were employed.

**Figure 2:**
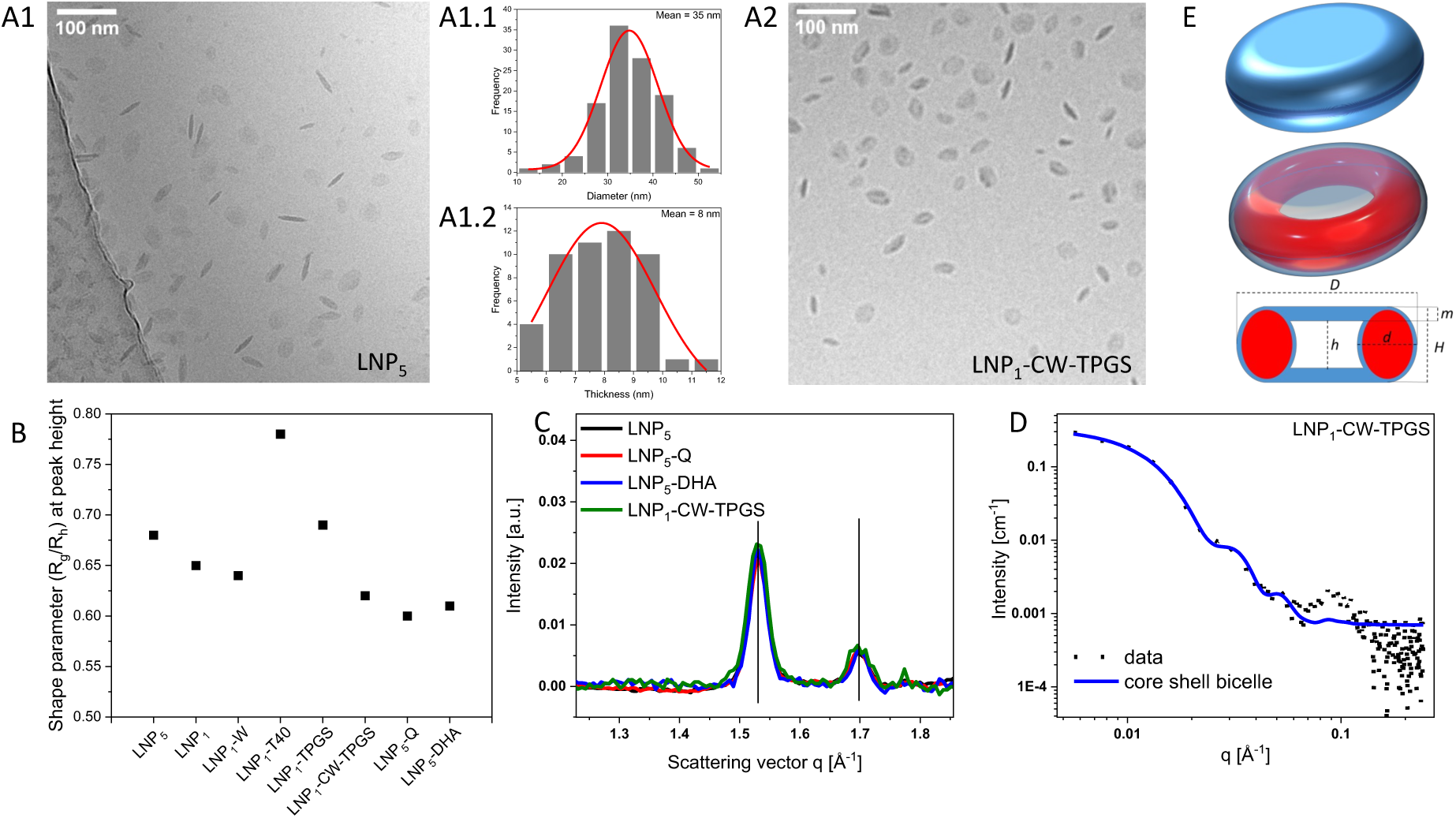
Shape and internal composition evaluation. (**A1**) Cryo-TEM image of LNP5 (Unloaded LNP with carnauba wax and red palm oil as lipid core, with both TPGS and Polysorbate 40 as surfactants), (**A1.1**) the average mean particle size distribution, and (**A1.2**) the average particle thickness distribution. (**A2**) Zoomed in cryo-TEM image of LNP1-CW-TPGS (Unloaded LNP with carnauba wax only lipid core and TPGS as surfactant) to provide a clearer view of the observed shape. (**B**) Shape parameter Rg/Rh determined at peak height for the various compositions of LNP formulation using AF4-MD shows values corresponding to spherical shapes. (**C**): WAXS plots of LNP5, LNP5-Q, LNP5-DHA and LNP1-CW-TPGS. (**D**) SAXS data and modelling by a core-shell bicelle for LNP1-CW-TPGS in water at a concentration of 11.02 mg.mL-1 based on lipid fraction. The geometry parameters correspond to the values described in the sketch under (E) with D = 37 nm; d = 9 nm; h = 10 nm; H = 17 nm; m = 3.5 nm. For the calculation, the SLD for water (core) and lipid (internal toroid) were applied. (**E**) Schematic representation of the particle shape corresponding to a disk-toroid hybrid based on structural insights gained from the experimental data using cryo-TEM and SAXS: the internal lipid toroid is stabilised by a monolayer of surfactant while the water core is stabilised by a double layer of surfactant at the interface between the core and aqueous particle environment.

The shape parameter (Rg/Rh), determined by AF4-MD at peak height (Figure 2 **B; Table S6**), spans 0.60 - 0.78 across formulations. Because these values fall below the hard-sphere limit (0.78), and far below the ∼ 2.0 expected for flat discs,^14–17^ they indicate complex, non-spherical architectures whose nuances require higher-resolution scattering methods.

To shed light onto internal structure and crystallinity wide-angle X-ray scattering (WAXS) was applied (Figure 2 **C**). Two reflections at q₁ = 1.53 Å⁻¹ and q₂ = 1.70 Å⁻¹ correspond to carnauba-wax spacings d₁ = 0.41 nm and d₂ = 0.37 nm.^18^ Drug loading with quinine or artemisinin leaves both lattice distances and crystallinity unchanged, suggesting the drugs reside outside the crystalline domains, possibly near the particle surface.

For deeper insight, SAXS data for LNP1-CW-TPGS (Figure 2 **D**) and LNP5 (**Figure S22**) at ∼ 11 mg ml⁻¹ (lipid basis) were modelled. Differences arise mainly between q = 0.025 and 0.05 Å⁻¹. A core-shell bicelle model, featuring distinct scattering length densities (SLDs) for core and shell, fits the data best (χ² < 1 × 10⁻⁵); while spherical and ellipsoidal models fail. A sharp peak at q = 0.09 Å⁻¹ remains unaccounted for by the bicelle model, but matches the signature of polysorbate-type micelles (∼ 7 nm) reported for Polysorbate-80 systems^12^ and noted in trace amounts by AF4 (see **Figures S4** and **S11**).

Fitted shell SLDs approximate that of carnauba wax (7.38 × 10⁻⁶ Å⁻²), whereas core SLDs drop from 9.15 × 10⁻⁶ Å⁻² (LNP1-CW-TPGS) to 8.19 × 10⁻⁶ Å⁻² (LNP5). Because water has SLD = 9.47 × 10⁻⁶ Å⁻², the reduced core SLD implies a water-rich interior - an almost empty core enclosed by a surfactant bilayer. This agrees with previous SAXS reports on solid LNPs that found similar SLD parity between water and surfactant layers.^12^ Modelled dimensions (Figure 2 **E**) align with the shape found by cryo-TEM but with different aspect ratios found: while 2:1 ratio was found using SAXS, cryo-TEM analysis leads to 4:1 ratio. This difference stems from the fact that SAXS is a batch method delivering an average size over the complete sample populations, while cryo-TEM analyses still a small fraction of the particles compared to the complete sample.

The observed distinctive biconcave geometry gives a larger surface-to-volume ratio than an equivalent sphere, favouring drug diffusion and membrane interactions.^19^ Differential scanning calorimetry of wax-oil matrices shows lowered enthalpy and onset temperatures, signatures of less-ordered α/β′ crystals,^10^ while the asymmetric TPGS/Polysorbate-40 shell promotes curvature. Together, these factors explain the torus-like SAXS patterns seen in wax nanoparticles^12^, and likely accelerate enzymatic access after injection, consistent with their rapid intracellular disassembly.

### Drug encapsulation – quantification and localisation

AF4-MD is uniquely suited to obtain, in a single run, conformation, size, molar mass, and composition distributions of heterogeneous nanoparticles across a broad size range.^14^ The channel’s semi-permeable membrane, here 10 kDa, defines the lower size limit: species smaller than the cut-off permeate the membrane and are swept into the cross-flow line instead of being separated in the channel. What might appear to be a drawback becomes an efficient way to quantify free drug in a drug-delivery system.^20, 21^ In the present work, we exploit that principle, determining the non-encapsulated fraction from a calibrated UV signal collected at the cross-flow outlet, while an in-line fluorescence detector pinpoints the drug associated with the LNP carrier.

**Figure 3** compares the two complementary approaches:

*(i) UV at the cross-flow outlet.* The cross-flow waste is routed directly to a UV-Vis detector. Any free drug that has diffused through the membrane appears as a UV peak. Calibration curves created from drug standards (Figures 3 **B1, B2**) relate peak area to concentration. A wavelength of 250 nm was selected because unloaded LNP5 shows no absorbance at this wavelength (Figure 3 **B3**), whereas the quinine-loaded sample (LNP5-Q) yields a clear signal. Integrating that peak and subtracting it from the total drug added during LNP formation gives an encapsulation efficiency of ∼90 %.
*(ii) In-line fluorescence.* Quinine and unloaded LNPs have different fluorescence maxima (450 nm and ∼350 nm, respectively). The quinine-loaded particles emit at wavelengths close to quinine (**Figure S24**). Batch measurements alone cannot reveal whether quinine is incorporated into the LNPs, but AF4-coupled fluorescence resolves the ambiguity. Overlapping elution profiles, higher fluorescence intensity at equal concentrations, and the red-shifted maximum prove that the drug co-elutes, and is therefore associated with the nanoparticles (Figure 3 **C1 and C2**).

**Figure 3:**
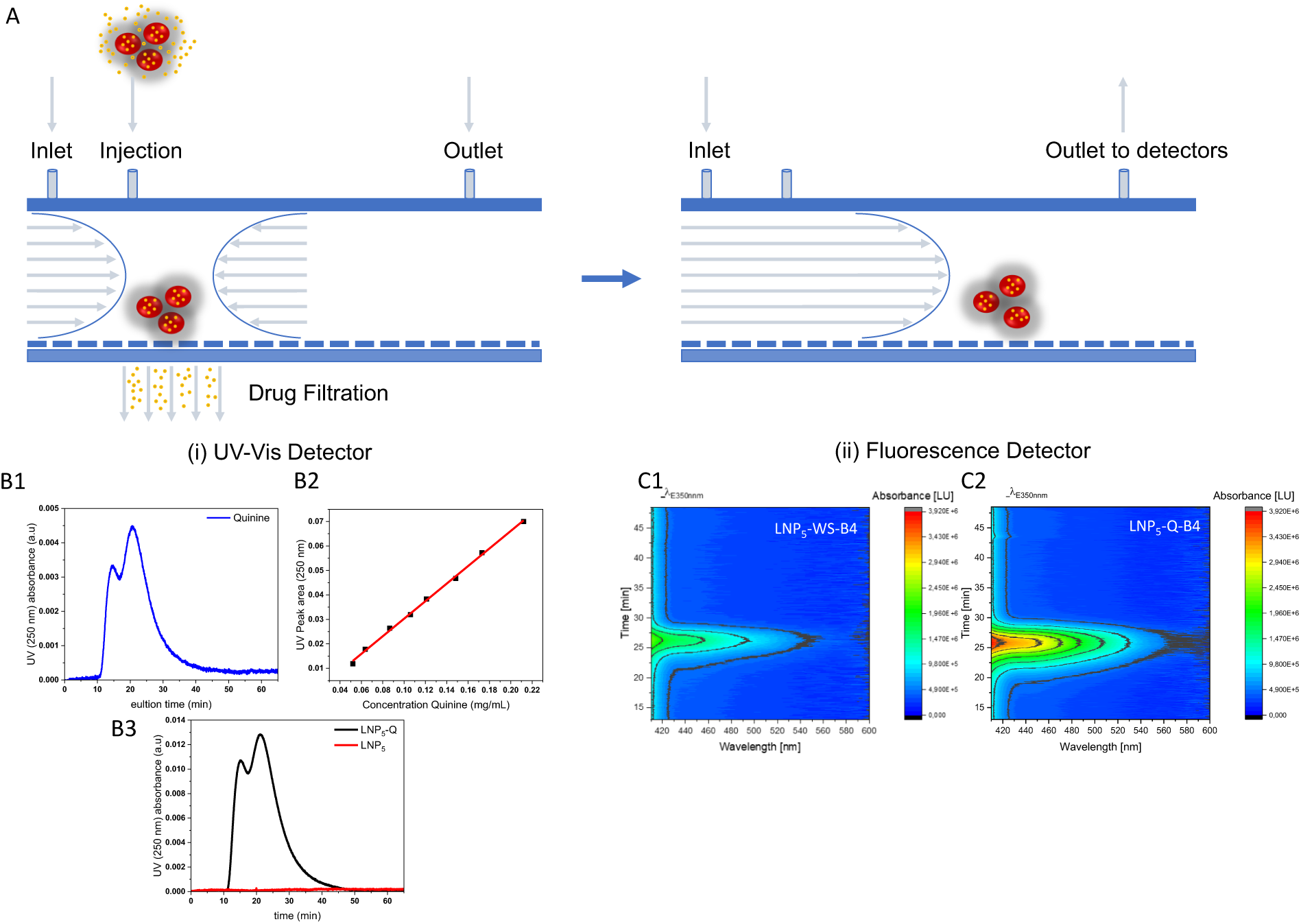
Drug encapsulation and quantification. (**A**) A schematic representation illustrating two alternative approaches for the indirect determination of the encapsulation efficiency of LNP, using AF4 coupled to either a UV-Vis detector or a fluorescence detector. A collection of small drug molecules filtered through the membrane of the AF4 channel is quantified using a pre-calibrated UV-Vis detector. The large drug-loaded particles separated along the channel are detected using the fluorescence detector. (**B1**) Elution profile showing the UV-Vis signal of quinine, with the cross-flow coupled to the UV-Vis detector. Wavelength set at 250 nm. (**B2**) The calibration curve is generated by injecting several concentrations of quinine, enabling the quantification of quinine present in the cross-flow waste. Wavelength set at 250 nm. (**B3**) Elution profile showing the UV-Vis signal of LNP5 and LNP5-Q, respectively. The cross-flow is coupled to the UV-Vis detector, set to a wavelength of 250 nm. **C1) and C2)** Fluorescence detection of the LNPs shows that the quinine is located in the particle due to increased intensity of the fluorescence signal at the same elution volume as the particle. The concentration of the particles is kept the same; thus, the increasing absorption intensity and broader absorption wavelength after excitation at 250 and 350 nm are the result of the quinine in the loaded LNP5-Q-B4 compared to the pure LNP5-WS-B4.

### Bioactivity in human cells

Intradermal or subcutaneous injection of LNPs is emerging as a promising approach for targeted local drug delivery, owing to their ability to provide sustained drug release over time. The biocompatibility of LNPs following skin injection appears to depend on the specific lipids and surfactants used in their formulation.^13^ To investigate this further, we assessed the cytotoxicity of a range of LNP variants on primary human fibroblasts and macrophages as examples of skin-resident tissue and immune cells that play key roles in skin homeostasis and repair. Fibroblasts produce the extracellular matrix (ECM) essential for skin structure, while macrophages function as immune sentinels and contribute to tissue regeneration. Their coordinated interaction is vital for maintaining skin integrity and responding to injury.^22, 23^ Co-culture of LNPs with fibroblasts and macrophages for 48 hours demonstrated very good biocompatibility up to a lipid concentration of 0.0024% for fibroblasts (Figure 4 **A1**) and up to 0.006% for macrophages (Figure 4 **A2**). This high level of biocompatibility was maintained even after 7 days of co-culture (**Figure S21 A** and **C**). At higher lipid concentrations, occasional morphological signs of cellular stress and apoptosis were observed (Figure 4 **B1** and **B2; Figure S21 B** and **D),** which corresponded to a slight but statistically significant reduction in cell viability. In contrast, the presence of surfactant and LNP payload had no measurable impact on cell viability (Figure 4 **A1 and A2; Figure S21 A and C**).

**Figure 4:**
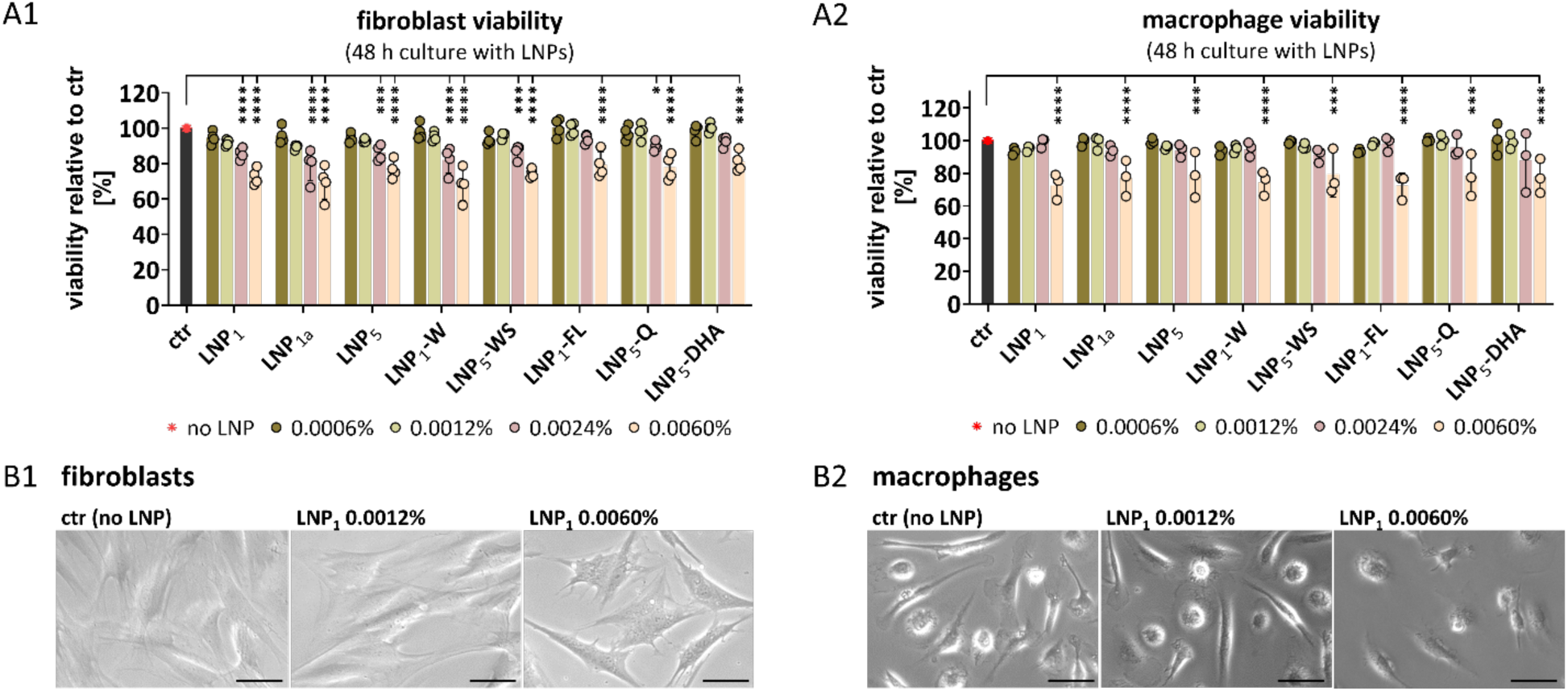
Cell viability after culture with LNP. **(A-B)** Monocyte-derived human macrophages and human dermal fibroblasts were cultured for 48 hours with different LNPs at different concentrations. LNP concentrations are indicated as % lipid fraction. LNP_1_, LPN_1a_, and LNP_5_ comprise different batches of LNP. LNP_1_-W are in pure water. LNP_5_-WS, the residual surfactants were removed. LNP_1_-FL, LNP_5_-Q, LNP_5_-DHA comprise LNP batches loaded with fluorescein (FL) or with the drugs quinine (Q) or dihydroartemisinin (DHA). **(A1)** Viability of fibroblasts was determined with the XTT assay. n=4 different fibroblast donors. **(A2)** Viability of macrophages was determined with the XTT assay. n=3 different macrophage donors. **(B1)** and **(B2)** Microscopic evaluation of macrophages and fibroblasts. Scale = 50 µm. **(A1/A2)** Two-way ANOVA with Tukey‘s multiple comparisons test. Significant differences compared to ctr (no LNP) are indicated. * p& 0.05, *** p& 0.001, **** p& 0.0001.

To further assess the cytocompatibility of LNPs, we investigated their impact on key fibroblast functions in tissue homeostasis, such as cell growth; and in tissue repair, such as extracellular matrix (ECM) production and myofibroblast activation,^24^ here simulated through stimulation with TGF-b. Consistent with viability assays, no effects on fibroblast proliferation were observed up to a lipid concentration of 0.0012%, whereas higher concentrations impaired proliferation (Figure 5 **A, Figure S22 A**). Neither the use of different LNP batches nor drug loading influenced fibroblast proliferation (**Figure S22 B**). Accordingly, LNPs at a lipid concentration of 0.0012% had no impact on fibroblast function, as evidenced by unchanged expression levels of key ECM components COL1A1, COL3A1, EDA-FN (fibronectin) and the myofibroblast marker ACTA2 (α-smooth muscle actin) (Figure 5 **B**).

**Figure 5:**
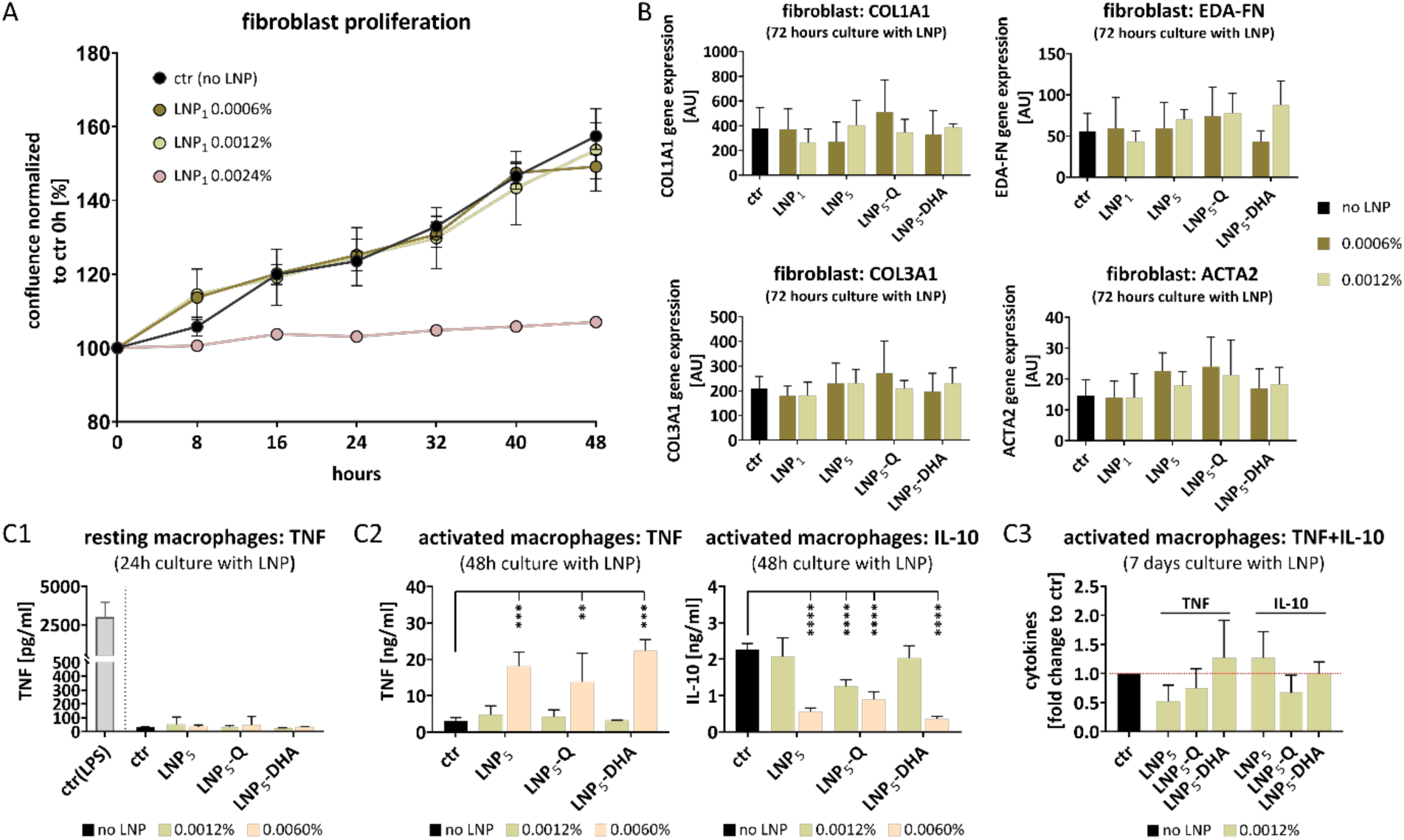
Effect of LNP on cell functions. **(A)** LNP_1_ at different concentrations were added to human dermal fibroblasts. Proliferation of fibroblasts was monitored via IncuCyte® live cell imaging. n=4. Two-way ANOVA with Dunnett‘s multiple comparisons test. Significant differences compared to ctr are indicated. ** p& 0.01, **** p& 0.0001. **(B)** LNPs at different concentrations were added to human dermal fibroblasts, which were then stimulated with TGF-b for 72 hours. Expression of matrix genes COL1A1, COL3A1, EDA-FN and myo-fibroblast activation gene ACTA2 were quantified. TGF-b-stimulated fibroblasts without LNP served as control. n=3. **(C)** Monocyte-derived human macrophages were cultured with LNPs at different concentrations. Culture without LNP served as a control. Macrophage activation was induced by stimulation with LPS 24 hours before the end of culture. **(C1)** TNF release of resting macrophages after 24 hours culture with LNPs. LPS-stimulated macrophages serve as a stimulation control (ctr-LPS). n=4. **(C2)** Release of TNF and IL-10 from activated macrophages after 48 hours culture with LNPs. n=4. **(C3)** Release of TNF and IL-10 from activated macrophages after 7 days culture with LNPs. Released cytokines are presented as fold change to control. n=4. **(B/C1/C2/C3)** Two-way ANOVA with Tukey‘s multiple comparisons test. Significant differences compared to ctr are indicated. * p& 0.05, ** p& 0.01, *** p& 0.001, **** p& 0.0001. **(A-E)** LNP concentrations are indicated as % lipid fraction. LNP_1_, LNP_5_, LNP_5_-Q, LNP_5_-DHA comprise different batches of LNP as outlined in **Figure 4**.

To evaluate the potential immunogenic effects that LNPs would induce after application in the skin,^25^ we analysed cytokine responses in resting macrophages after LNP exposure. No induction of TNF-α - a representative pro-inflammatory cytokine macrophages produce in response to their activation,^26^ for example, with LPS - was detected after 48 hours of co-culture (Figure 5 **C1**), nor after 7 days (data not shown). We further examined whether LNPs modulate inflammatory responses in pre-activated macrophages and exemplarily analysed the release of TNF as an inflammatory macrophage signal, and the release of IL-10 as a macrophage-derived signal for inflammatory resolution.^26, 27^ Consistent with viability data, LNP concentrations up to 0.006% had no effect on cytokine release after 48 hours or 7 days of co-culture (Figure 5 **C2, C3**). However, for LNPs with 0.006% lipid concentrations, a marked increase in TNF-α production, along with a decrease in the anti-inflammatory cytokine IL-10, was observed (Figure 5 **C2**), suggesting a concentration-dependent pro-inflammatory shift.

### Biocompatibility in the skin

Next, we assessed the behaviour of lipid nanoparticles (LNPs) under more native-like conditions by employing a human *ex vivo* skin culture model (**Figure S23 A**) to investigate LNP biocompatibility in a biologically and physiologically relevant context.^28^ The skin maintained structural integrity over a 14-day culture period, with only minor epidermal detachment observed by day 14 (**Figure S23 B**). Notably, the dermis and hypodermis remained well-preserved, showing minimal and consistent levels of apoptosis throughout the culture period (**Figure S23 C**). In summary, our *ex vivo* skin culture model maintains good tissue integrity for up to 14 days, consistent with findings from previous studies,^29, 30^ and provides a suitable platform for investigating potential adverse effects of LNPs within this timeframe.

LNPs at various lipid concentrations were injected at the dermis-hypodermis interface, as illustrated in Figure 6 **A**. Histological analysis of skin samples four days post-injection revealed well-maintained skin architecture. No differences were observed between untreated skin, PBS-injected controls, and skin injected with LNPs at 0.024% lipid concentration (Figure 6 **B**). However, skin injected with LNPs at 1.2% lipid concentration showed mild disruption in adipocyte integrity, suggesting lipolysis. Lipolysis is a key metabolic process in which lipids stored in fat cells are broken down into glycerol and fatty acids, typically in response to fat activation and stress.^31–33^

**Figure 6:**
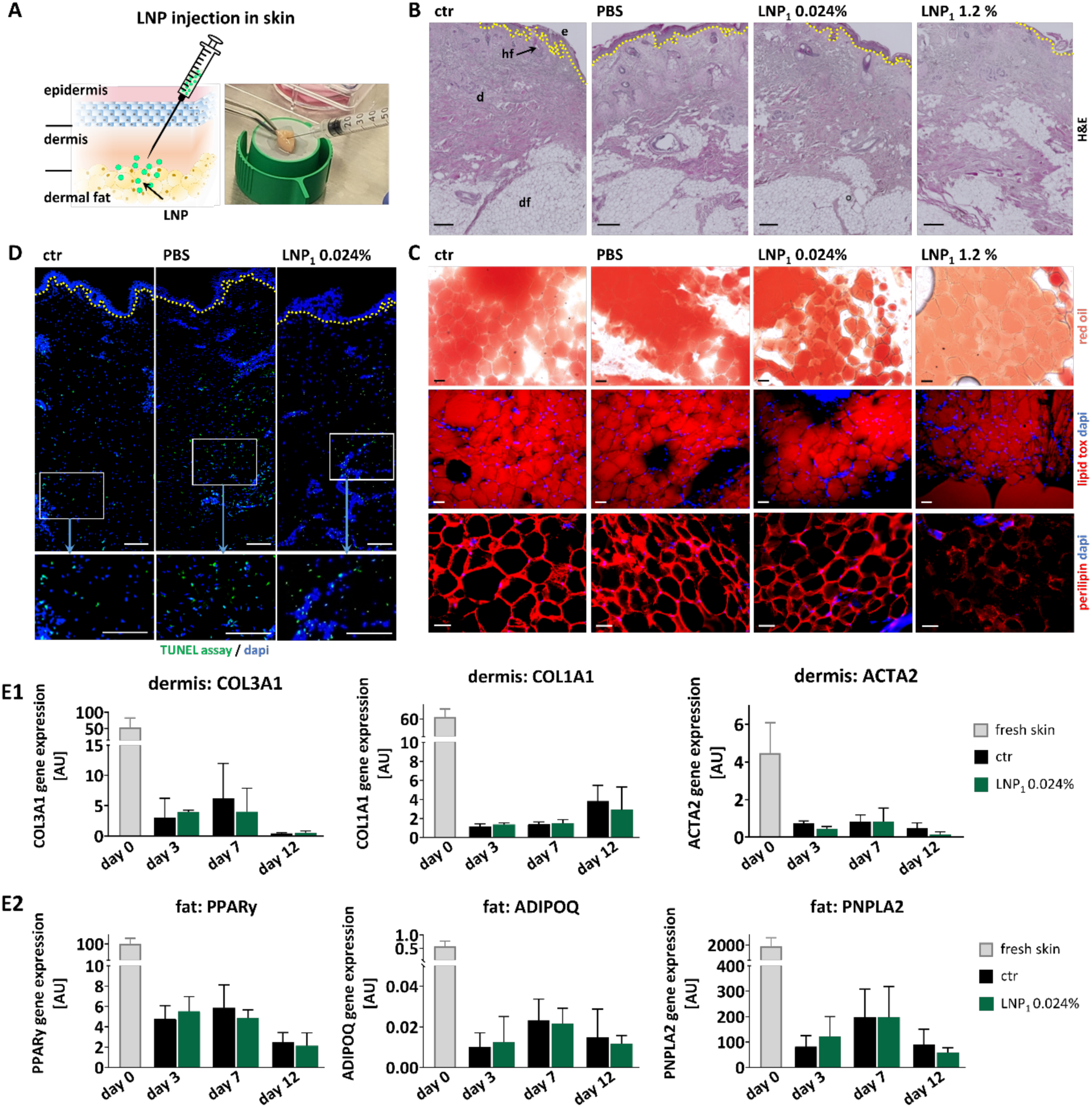
Biocompatibility of LNP in human skin. **(A)** Illustration of LNP injection into human skin. **(B-E)** 100µl of LNP_1_ at 1.2 % lipid fraction (LNP_1_ 1.2 %) or of LNP_1_ diluted in PBS at 0.024% lipid fraction (LNP_1_ 0.024 %) were injected in the deeper dermis/dermal fat area of human skin. Untreated human skin served as a control (ctr). Injection of 100µl PBS served as a treatment control (PBS). Skin samples were cultured ex vivo as shown in the Supplementary Figure 23A**. (B)** Hematoxylin and Eosin (H&E) staining of histological skin sections. Skin culture ex vivo for four days. scale = 300µm **(C)** Evaluation of lipolysis in the dermal fat by staining with red oil, lipid tox and perilipin in histological sections. Skin culture ex vivo for four days. scale = 50µm. **(D)** Evaluation of skin viability via TUNEL staining of histological sections. Skin culture ex vivo for two days. scale = 200µm. **(E1)** and **(E2)** Gene expression analysis of dermis and dermal fat isolated from different skin samples after culture ex vivo for three days, 7 days and 12 days. Statistical analysis was performed using two-way ANOVA followed by Tukey’s multiple comparisons test. No statistically significant differences were observed between the groups. n=4

The occurrence of lipolysis in skin injected with LNPs at 1.2% lipid concentration was supported by lipid staining (Red Oil and LipidTOX) and the observed loss of perilipin, a key lipid droplet-associated protein^34^ (Figure 6 **C**). In contrast, adipocyte morphology remained intact in all other conditions. TUNEL assay confirmed good viability of skin injected with LNPs at 0.024%, with apoptosis levels comparable to PBS-treated controls (Figure 6 **D**). To assess potential functional changes in skin cells following LNP injection, dermal and fat layers were separated post-culture for gene expression analysis. Although cultured skin showed an overall decrease in gene expression compared to fresh skin, no significant differences were observed between control and LNP-injected skin (0.024%) in expression of extracellular matrix-related genes (COL1A1, COL3A1, ACTA2) in the dermis (Figure 6 **D**). Similarly, genes related to adipogenesis, adipocyte activation and lipolysis (PPARγ, ADIPOQ, PNPLA2, respectively)^35–37^ were not differentially expressed between control and LNP-treated fat tissue (Figure 6 **D**).

In summary, the evaluation of LNPs in the human *ex vivo* skin model demonstrated excellent biocompatibility at lipid concentrations up to 0.024%, with no structural or functional impairments observed at this dose.

Solid lipid nanoparticles are a major subclass of lipid-based nanocarriers, particularly well-suited for targeted drug delivery.^38^ Effective drug delivery through the skin requires efficient penetration across its layers, which has been suggested to depend on specific physicochemical properties of the LNPs, such as solvent polarity and particle size.^39^ To assess whether LNPs can achieve skin penetration, we injected fluorescein-loaded LNPs at a lipid concentration of 0.024% into our human ex vivo skin model. The injected skin was placed in a transwell system with a membrane pore size that permits the passage of LNPs but not cells (Figure 7 **A**). Supernatants were collected at defined time points following injection, and fluorescence intensity was measured using a plate reader. At each time point, the culture medium was fully replaced to ensure accurate quantification of continuous release. We observed already one hour after injection a fluorescence signal in the supernatant, which peaked at 24 hours, followed by a gradual decline over subsequent time points (Figure 7 **B**). It is noteworthy that the measured total fluorescence corresponds to a maximum lipid concentration of approximately 0.002% (Figure 7 **C**). These findings indicate that a small but detectable fraction of LNPs was able to actively pass through the dermal and hypodermal layers.

**Figure 7.**
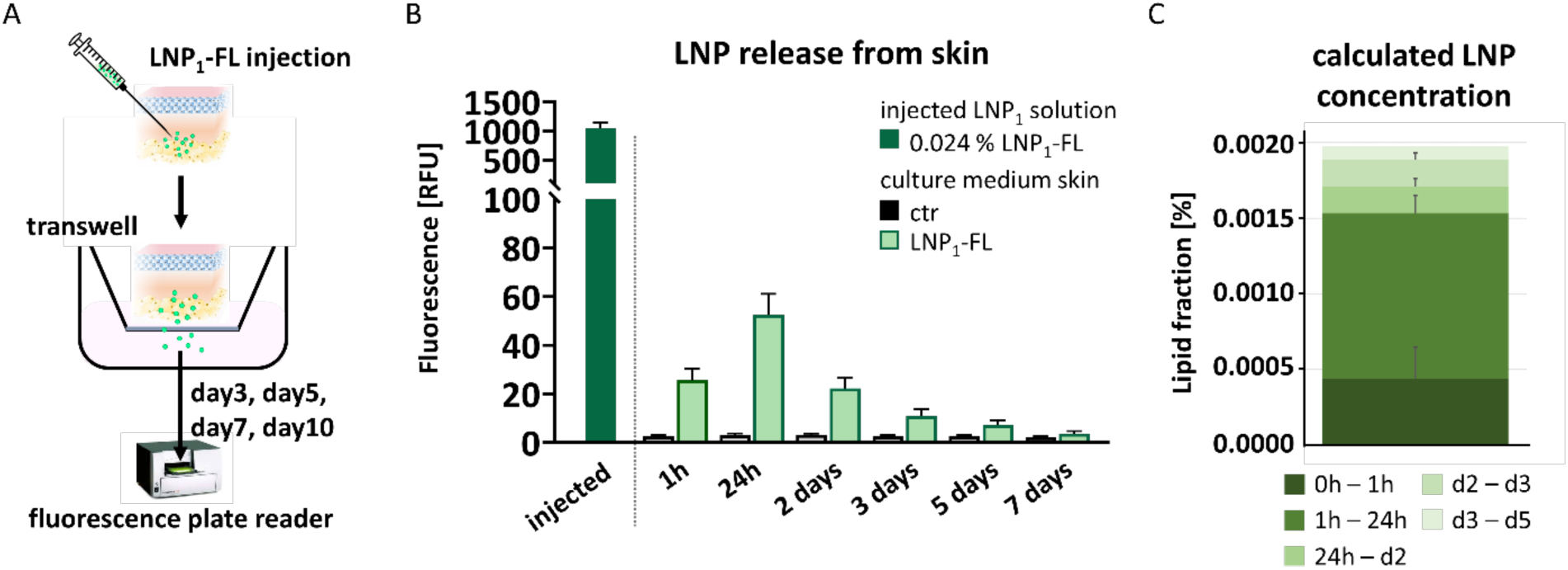
Penetration of LNP through the dermal and fat layer of human skin. **(A)** Illustration of experimental set-up. **(B/C)** 100µl of LNP_1_ loaded with fluorescein (LNP_1_-FL) diluted at 0.024% lipid fraction was injected in the deeper dermis/dermal fat area of human skin. Untreated human skin served as control (ctr). Skin samples were placed in a transwell (0.4 µm pore size) in a culture plate and cultured ex vivo. After the indicated time points, the culture medium was replaced with fresh medium and subjected to fluorescence analysis. **(B)** Quantification of green fluorescence of collected culture medium and initially injected LNP_1_-FL solution. n=4. Statistical analysis was performed using two-way ANOVA Šídák’s multiple comparisons test. ** p& 0.01, **** p& 0.0001. **(C)** Calculated lipid fraction that accumulated in the medium during the culture periods of the skin injected with LNP_1_-FL. n=4.

In summary, our data suggest that low levels of LNPs can penetrate through the deeper layers of the skin in this model, supporting their potential for transdermal drug delivery. Our findings are supported by a previous study showing that lipid nanoparticles smaller than 100 nm fall within the optimal size range for deep skin penetration.^40^

## CONCLUSIONS

This study delineates a clear design space for solid-liquid LNP depots and demonstrates that rational matrix and surfactant engineering yields carnauba-wax/red-palm-oil lipid nanoparticles (LNPs) that combine long-term colloidal stability with high drug-loading capacity and excellent tissue compatibility.

Dispersions prepared by phase inversion remain monodisperse for at least 18 months at 4 °C, retaining a hydrodynamic diameter below 50 nm and a polydispersity index ≤ 0.20. The mixed TPGS/polysorbate-40 corona and inclusion of red-palm oil are both critical: removing either component broadens the size distribution and induces bimodal populations. Quinine and dihydroartemisinin can be encapsulated at ≈ 90 % efficiency without altering particle size or wax lattice order. Zeta potentials of −8 to −15 mV prevent aggregation even at lipid concentrations exceeding 10 mg mL⁻¹, enabling highly concentrated injectable depots.

Dominant distinctive disc-toroid popukation that provides an enlarged surface-to-volume ratio has been confirmed complementary by Cryo-TEM and SAXS. The hybrid shape is not just a morphological curiosity: variation in surface area dictate how much membrane each particle can contact and therefore influence cellular uptake. Particles with disc-toroid geometry are expected to interact with cell membranes differently from their spherical counterparts, potentially modulating internalisation kinetics. Systematic uptake studies in keratinocytes, fibroblasts and macrophages are thus a logical next step to correlate shape with biological fate.

Cytotoxicity thresholds lie well above therapeutically relevant lipid doses (≤ 0.006 % for macrophages and ≤ 0.0024 % for fibroblasts), and ex-vivo human skin retains normal architecture and gene expression after injection of 0.024 % lipid. Only trace amounts (≤ 0.002 %) traverse the dermis within 24 h, confirming depot-like behaviour.

Collectively, these insights converge on a formulation window in which disc-toroid LNPs combine long-term physical stability, high drug payload and excellent tissue compatibility. They therefore constitute a versatile platform for minimally invasive delivery of small-molecule or biopharmaceutical therapeutics. Future work should couple this structural blueprint with in-vivo pharmacokinetics and ligand-directed surfactant coronas to fully exploit their therapeutic potential.

## MATERIALS AND METHODS

### Materials

Phosphate buffer saline (PBS) tablets (Sigma-Aldrich); sodium azide (NaN3) (Sigma-Aldrich), sodium chloride (NaCl) (Merck), MilliQ ultrapure water, carnauba wax (Sigma-Aldrich), red palm oil concentrate (Sigma-Aldrich), d-a-Tocopheryl polyethylene glycol 1000 succinate (TPGS) (Sigma-Aldrich) and polysorbate 40 (Sigma-Aldrich). Quinine (Sigma-Aldrich) and dihydroartemisinin (Apidogen) were used as the payload. All materials were used as received unless otherwise stated.

### Preparation of unloaded and loaded lipid nanoparticles

For the preparation of dispersion of the lipid nanoparticles (LNP), carnauba wax, red palm oil concentrate, d-a-Tocopheryl polyethylene glycol 1000 succinate (TPGS), and polysorbate 40 are mixed (**Table 1**). The mixture is heated up to 90°C ± 2°C to melt and stirred (500 rpm) until a homogeneous, clear mixture is obtained. The needed amount of drug is added to the lipid mixture and is stirred (500 rpm) until it fully dissolves. The needed amount of deionised water with the NaCl dissolved in it is heated up to 90°C ± 2°C, and it is added dropwise to the homogeneous mixture obtained under stirring. The obtained dispersion is cooled down under stirring (300 rpm) to 20°C ± 2°C to give the nanoparticle dispersion.

**Table 1:** The composition of the LNPs containing a drug.

| Compounds | Amount in w/w parts |
| --- | --- |
| Carnauba wax | 1.00 to 5.00 |
| Red palm oil concentrate (30% tocotrienols) | 0.10 to 0.50 |
| d-α-Tocopheryl polyethylene glycol 1000 succinate (TPGS) | 0.70 to 3.50 |
| Polysorbate 40 | 0.50 to 2.50 |
| Drug<sup>a</sup> | q.s. |
| NaCl | 0.9 |
| Water | up to 100.0 |

<sup>a</sup> Drug (Quinine or Dihydroartemisinin); q.s. quantum sufficient

### Purification and dialysis of samples

(a) The procedure for purification of the prepared LNPs involves the removal of the residual amount of surfactant used in the preparation of LNPs and is described as “washed from surfactants” in **Table 2**. A volume of 20 ml of the tested dispersion was filtered through a 0.45 µm glass fibre syringe filter and introduced into a Spectrum™ Spectra/Por™ 7 dialysis tubing with a 50 kD MWCO. A permeable membrane allowed surfactants to pass through, while restricting particle movement. It was sealed at both ends and immersed in a 5-liter glass beaker with 4500 ml of 0.9% NaCl solution in MilliQ ultrapure water at 25°C for 48 hours with continuous stirring, replacing the washing solution every 12 hours. After 48 hours, ten 5-ml samples of the washing solution were scanned in the UV region from 200 nm to 400 nm, using 0.9% NaCl in MilliQ ultrapure water as a blank. The signal was analysed to determine the lack of measurable value (XL) of solutes in the final washing solution. The calculation of XL is expressed by the equation:

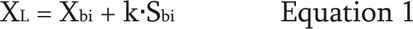

**Table 2:** Lipid nanoparticle detailed description.

| Sample | Description |
| --- | --- |
| LNP<sub>5</sub> | Unloaded lipid nanoparticles dispersed in 0.9% NaCl solution, 5.5% lipid fraction, with carnauba wax and red palm oil as lipid core, with both TPGS and Polysorbate 40 as surfactants. |
| LNP<sub>5</sub>-WS | Unloaded lipid nanoparticles dispersed in 0.9% NaCl solution, 5.5% lipid fraction, with carnauba wax and red palm oil as lipid core, with both TPGS and Polysorbate 40 as surfactants.<br>Washed from residual surfactant |
| LNP<sub>5</sub>-Q | Quinine (Q)-loaded lipid nanoparticles dispersed in 0.9% NaCl solution, 5.5% lipid fraction, with carnauba wax and red palm oil as lipid core, with both TPGS and Polysorbate 40 as surfactants.<br>10% (w/w) quinine in lipid |
| LNP<sub>5</sub>-DHA | Dihydroartemisinin (DHA)-loaded lipid nanoparticles dispersed in 0.9% NaCl solution, 5.5% lipid fraction, with carnauba wax and red palm oil as lipid core, with both TPGS and Polysorbate 40 as surfactants.<br>3 % (w/w) DHA in lipid |
| LNP<sub>1</sub> | Unloaded lipid nanoparticles dispersed in 0.9% NaCl solution, 1.1% lipid fraction, with carnauba wax and red palm oil as lipid core, with both TPGS and Polysorbate 40 as surfactants. |
| LNP<sub>1</sub>-W | Unloaded lipid nanoparticles dispersed in ultrapure water, 1.1% lipid fraction, with carnauba wax and red palm oil as lipid core, with both TPGS and Polysorbate 40 as surfactants. |
| LNP<sub>1</sub>-Q | Quinine (Q)-loaded lipid nanoparticles dispersed in 0.9% NaCl solution, 1.1% lipid fraction, with carnauba wax and red palm oil as lipid core, with both TPGS and Polysorbate 40 as surfactants.<br>10% (w/w) quinine in lipid |
| LNP<sub>1a</sub> | Unloaded lipid nanoparticles dispersed in 0.9% NaCl solution, 1.2% lipid fraction, with carnauba wax and red palm oil as lipid core, with both TPGS and Polysorbate 40 as surfactants. |
| LNP<sub>1</sub>-FL | Fluorescein-loaded lipid nanoparticles dispersed in 0.9% NaCl solution, 1.1% lipid fraction, with carnauba wax and red palm oil as lipid core, with both TPGS and Polysorbate 40 as surfactants<br>0.45% (w/w) fluorescein in lipid |
| LNP<sub>1</sub>-CW-TPGS | Lipid nanoparticles dispersed in ultrapure water, 1.1% lipid fraction, with carnauba wax as lipid core, with TPGS as surfactant. |
| LNP<sub>1</sub>-CW-T40 | Lipid nanoparticles dispersed in ultrapure water, 1.1% lipid fraction, with carnauba wax as lipid core, with Polysorbate 40 as surfactant |
| LNP<sub>1</sub>-TPGS | Unloaded lipid nanoparticles dispersed in ultrapure water, 1.1% lipid fraction, with carnauba wax and red palm oil as lipid core, with TPGS as surfactant. |
| LNP<sub>1</sub>-T40 | Unloaded lipid nanoparticles dispersed in ultrapure water, 1.1% lipid fraction, with carnauba wax and red palm oil as lipid core, with Polysorbate 40 as surfactant. |
| LNP<sub>5</sub>-B2 | Unloaded lipid nanoparticles dispersed in 0.9% NaCl solution, 5.5% lipid fraction, with carnauba wax and red palm oil as lipid core, with both TPGS and Polysorbate 40 as surfactants. |
| LNP<sub>5</sub>-Q-B2 | Quinine (Q)-loaded lipid nanoparticles dispersed in 0.9% NaCl solution, 5.5% lipid fraction, with carnauba wax and red palm oil as lipid core, with both TPGS and Polysorbate 40 as surfactants.<br>10 % (w/w) quinine in lipid |
| LNP<sub>5</sub>-DHA-B2 | Dihydroartemisinin (DHA)-loaded lipid nanoparticles dispersed in 0.9% NaCl solution, 5.5% lipid fraction, with carnauba wax and red palm oil as lipid core, with both TPGS and Polysorbate 40 as surfactants.<br>5 % (w/w) DHA in lipid |
| LNP<sub>5</sub>-B3 | Unloaded lipid nanoparticles dispersed in 0.9% NaCl solution, 5.5% lipid fraction, with carnauba wax and red palm oil as lipid core, with both TPGS and Polysorbate 40 as surfactants. |
| LNP<sub>5</sub>-B4 | Unloaded lipid nanoparticles dispersed in 0.9% NaCl solution, 5.5% lipid fraction, with carnauba wax and red palm oil as lipid core, with both TPGS and Polysorbate 40 as surfactants. |
| LNP<sub>5</sub>-WS-B4 | Unloaded lipid nanoparticles dispersed in 0.9% NaCl solution, 5.5% lipid fraction, with carnauba wax and red palm oil as lipid core, with both TPGS and Polysorbate 40 as surfactants.<br>Washed from residual surfactant |
| LNP<sub>5</sub>-Q-B4 | Quinine (Q)-loaded lipid nanoparticles dispersed in 0.9% NaCl solution, 5.5% lipid fraction, with carnauba wax and red palm oil as lipid core, with both TPGS and Polysorbate 40 as surfactants.<br>20 % (w/w) quinine in lipid |
| LNP<sub>5</sub>-DHA-B4 | Dihydroartemisinin (DHA)-loaded lipid nanoparticles dispersed in 0.9% NaCl solution, 5.5% lipid fraction, with carnauba wax and red palm oil as lipid core, with both TPGS and Polysorbate 40 as surfactants<br>3 % [w/w] DHA in lipid |
| LNP | Unloaded lipid nanoparticles dispersed in 0.9% NaCl solution, 0.6% lipid fraction, with carnauba wax and red palm oil as lipid core, with both TPGS and Polysorbate 40 as surfactants |
| LNP-Q | Quinine (Q)-loaded lipid nanoparticles dispersed in 0.9% NaCl solution, 1.2% lipid fraction, with carnauba wax and red palm oil as lipid core, with both TPGS and Polysorbate 40 as surfactants. 8 % (w/w) quinine in lipid |

With Xbi, the mean of the blank measurements, which serves as a baseline, Sbi is the standard deviation of the blank measurements, indicating measurement variability; k is a numerical factor chosen according to the desired confidence level for detection. In our tests k = 3. The equation sets a limit of detection, ensuring reliable analytical results and aiding decision-making for an extra 12-hour washing period(s) based on analyte quantities in the samples. Dialysis of the samples continued until the absence of an absorbance peak corresponding to the surfactant used. All samples prepared were packed in aseptic conditions in a Class 2A biological safety cabinet (Biobase Class IIA Biological Safety Cabinet).

(b) Dialysis was carried out against MilliQ water for 24 hours, replacing the washing solution every 6 hours (Spectra/Por® Float-A-Lyzer® G2 Volumen 1 ml, 8-10 kDa MWCO membrane) at 200 rpm. The dialysis samples were used for batch-mode dynamic light scattering (DLS) and small-angle x-ray scattering (SAXS).

### Characterisation of lipid nanoparticles Dynamic light scattering (DLS)

Batch-mode DLS studies were carried out on a Zetasizer Nanoseries (Malvern Instruments, UK), equipped with Zetasizer software (version 8.02). Data were collected at a scattering angle of 173° and with a laser wavelength of λ = 632 nm. Data evaluation was carried out by using Zetasizer Software (version 8.02). Measurements were performed in an aqueous solution of 0.9% NaCl or MilliQ water using disposable cuvettes (ZEN0040). The automatic attenuator factor was between 6 and 7. Measurements were performed at 25°C, all samples were equilibrated for 5 minutes before each measurement, and the software automatically selected the acquisition time for each measurement according to the sample conditions. The mean particle size and size distribution were assessed by intensity (intensity-PSD). All measurements were performed in a minimum of triplicate.

### Zeta potential

The zeta potential studies were carried out on a Zetasizer Nanoseries (Malvern Instruments, UK), equipped with Zetasizer software (version 8.02). Data evaluation was carried out by using Zetasizer Software (version 8.02). Measurements were performed in an aqueous solution of 1 mM NaCl using disposable folded capillary cells (DTS1070). Measurements were performed at 25°C, and all samples were equilibrated for 5 minutes before each measurement. The results were determined by the mean of five measurements. Between each measurement, there was a 300-second delay. The model for the F(ka) selection was Smoluchowski (1.5). The measurement setting was selected with automatic attention and automatic voltage. The analysis mode was monomodal unless otherwise stated.

### Asymmetric flow field-flow fractionation (AF4)

AF4 measurements were performed with an Eclipse Neon Asymmetric flow field-flow fractionation (AF4) system (Wyatt Technologies Corp., Germany). The system is equipped with a DAWN Neon MALLS (18 angles, operating at a wavelength of 659 nm) with online QELS, OptiLab dRI (operating at a wavelength of 658 nm) (Wyatt Technologies Corp., Germany), and Agilent 1260 Infinity II MWD, an Agilent Infinity II isocratic pump with an integrated degassing system and an autosampler (Agilent Technologies 1260 Infinity series, Agilent Technologies, USA) (AF4-UV-QELS-MALS-dRI).

Measurements were carried out in a carrier liquid of 10 mM PBS buffer at pH 7.4, containing 200 mg.L^-1^ NaN3. An Eclipse short channel with DCM and a defined channel height of 350 µm asymmetric channel spacer was connected. A regenerated cellulose (RC) ultrafiltration membrane (Wyatt Technologies Corp., Germany) with a molecular weight cut-off (MWCO) of 10 kDa was used. The samples were injected into the AF4 separation channel using the autosampler. The AF4 method was optimised to achieve optimal fractionation of the lipid nanoparticles (see Supplementary **Figure S1** and **Table S5**). The data was collected by Astra 8.1.2 software (Wyatt Technologies Corp., Germany). The following method was applied: Detector flow rate was set to 0.5 ml.min^-1^, for the separation of an isocratic step with a Vx of 3 ml.min^-1^ for 9 min (Elution; Focus; Focus Inject; Elution), followed by an exponential Vx gradient (slope 8) from 3 to 0.05 ml.min^-1^ within 28 min was used. The last steps consisted of an isocratic Vx of 0.05 ml.min^-1^ for 7 min, followed by an isocratic Vx of 0 ml.min^-1^ for 15 min. The MWD lamp was set to 250 nm, 280 nm, 300 nm, 310 nm, and 330 nm. Three injections of 25 µl and one of 250 µL were performed for each sample, and the injection flow rate was set to 0.2 ml.min^-1^. Recovery tests were performed for each sample to determine that no adsorption phenomena took place at the membrane interface. For the collection of time-dependent fluorescence spectra, a FLD Infinity II 1260 (Agilent Technologies) was included in the setup. The excitation wavelengths were set to 250 and 350 nm. The emission spectra were recorded from 250 – 600 nm during a time of 10 - 40 min of elution using the separation protocol as described above.

### Fluorescence in batch

Fluorescence spectra were recorded using an FS5 Spectrofluorometer (Edinburgh Instruments Corp., USA), equipped with a xenon flash lamp. LNP samples were prepared by dispersing 10 μL of the LNP suspension in 990 μL of 0.9% NaCl. Excitation was set at 288 nm, and emission spectra were collected from 300 – 600 nm. The slit widths were set to 3 nm. Measurements were performed at room temperature.

### Transmission Electron Microscopy (TEM)

Cryo-transmission electron microscopy (cryo-TEM) images were recorded in Libra 120 microscope (Carl Zeiss Microscopy Deutschland GmbH, Oberkochen, Germany). 2 µL to 4 µL of specimen was placed onto a holey carbon TEM grid (Quantifoil R3.5/1, 300 mesh), blotted with filter paper for 0.2 s to 1 s and vitrified in liquid ethane at -178 °C using a Grid Plunger (Leica Microsystems GmbH, Wetzlar, Germany). Frozen grids were transferred into Gatan 626 (Gatan GmbH, München, Germany) cryo-TEM holder. Images were recorded at an accelerating voltage of 120 kV while keeping the specimen at -170 °C. To study the size distribution of particles, ImageJ software (version) was used. To quantify the size distribution, 30 – 40 spherical particles and 10 – 20 elongated particles were manually measured in ImageJ, from three different images to obtain the average size distribution. The raw data were processed in Origin Pro (version) and plotted using a Gaussian distribution curve.

### X-ray Scattering (SAXS and WAXS)

Small-angle X-ray scattering (SAXS) and wide-angle X-ray scattering (WAXS) measurements were performed on a Ganesha 300 XL+ (SAXLAB) in vacuum. The instrument is equipped with a pinhole-focused X-ray beam (CuK*a*, *l*=1.542 Å) and a Pilatus 300k pixel detector (pixel size: 172 x 172 µm^2^). SAXS and WAXS data were recorded at a sample detector distance of 1052 mm and 110 mm, respectively. Quartz glass capillaries were loaded with lipid nanoparticle solutions at different concentrations (based on lipid fraction) of 11, 4, 2, 0.5 mg.ml^-1^ in pure water or 0.9 % NaCl. SAXS integration times were chosen between 2-4 h, while WAXS data was only obtained for the 11 mg.ml^-1^ concentration, integrating for 30 minutes for each sample. 2D data were azimuthally averaged to obtain 1D scattering curves, after standard data correction procedures (e.g. dark current subtraction, flat field correction). SAXS data were normalised and converted to absolute intensity using water as a standard and then referenced to the respective pure solution without sample. SAXS data for LNP that consist only of a carnauba wax core (LNP1-CW-TPGS) and LNP5 in water were modelled in SasView vers. 5.0.5^41^using the core-shell bicelle model. X-ray CuKα scattering length densities (SLD) of the carnauba wax (C7H5HgNO3) and water were calculated using the NIST tool^42^ to SLDcarnauba = 7.38 · 10^!"^ Å^-2^ and SLDH2O = 9.47 · 10^!"^Å^-2^, respectively. WAXS data was scaled to the same background intensity level (amorphous SiO2 background) and then corrected for the background using a spline function.

### Biocompatibility studies Human samples

Human skin samples were obtained from clinically healthy donors using surplus tissue collected during elective surgical procedures (breast and abdominal areas) at the Department of Orthopaedics, Trauma Surgery, and Plastic Surgery, University Hospital Leipzig. In addition, peripheral blood samples were collected from healthy adult volunteers. All human samples were collected following written informed consent and with prior approval from the Ethics Committee of Leipzig University (approval number EK092-20), in accordance with the principles of the Declaration of Helsinki.

### Isolation and culture of primary human macrophages and fibroblasts

Human macrophages were generated as previously described.^43^ Briefly, peripheral blood mononuclear cells (PBMCs) were isolated from human blood by density-gradient centrifugation using Ficoll-Paque PLUS (Cytiva Sweden AB). Monocytes were then enriched from PBMCs to a purity of >95% using the CD14+ Cell Isolation Kit (Miltenyi). Enriched monocytes were seeded at a density of 4 × 10⁵ cells/ml in RPMI-1640 medium (Anprotec) supplemented with 1% (v/v) PenStrep (Anprotec), as well as 10% (v/v) heat-inactivated fetal calf serum (FCS, Anprotec) to complete RPMI medium. Monocyte-to-macrophage differentiation was induced by the addition of 50 ng/ml macrophage colony-stimulating factor (M-CSF, Miltenyi) and continued culture for six days under standard culture conditions.

Primary dermal fibroblasts were isolated from human skin tissue as previously described.^43^ Briefly, epidermal layers were separated from the dermis using Dispase II (Roche Diagnostics), followed by enzymatic digestion of the dermal compartment with collagenase (Sigma-Aldrich) to release fibroblasts. The resulting cell suspension was filtered through a 70 µm cell strainer (Greiner Bio-One) to eliminate residual tissue fragments. Fibroblasts were cultured in Dulbecco’s Modified Eagle Medium (DMEM; Anprotec) supplemented with 1% (v/v) PenStrep (Anprotec) and 10% (v/v) FCS to complete DMEM. Fibroblasts were grown to 80 % confluence, and only cells between passages 2 and 4 were used for downstream experiments.

Macrophages and fibroblasts were maintained at 37 °C in a humidified atmosphere containing 5% CO₂ for the duration of all experiments.

### LNP Stimulation of Macrophages and Fibroblasts

Macrophages were seeded at a density of 80,000 cells per well in 48-well plates using 250 µl of complete RPMI-1640 medium. Cells were then stimulated with 250 µl of various LNP formulations, prepared at the indicated concentrations in R10. For macrophage activation, cells were additionally stimulated with 100 ng/ml lipopolysaccharide (LPS, Sigma), as described previously.^44^ Fibroblasts were seeded at 20,000 cells per well in 24-well plates in 500 µl of complete DMEM and stimulated with 500 µl of LNP formulations prepared at different concentrations in D10. For cell viability assays, macrophages and fibroblasts were seeded in 96-well plates at 15,000 and 2,000 cells per well, respectively, in 100 µl of their respective complete media. After cell attachment, 100 µl of the corresponding LNP formulations were added to each well.

### *Ex vivo* human skin model

The *ex vivo* skin culture model was established based on the method described by Wilkinson *et al.*^45^ Human skin samples were obtained from surgical procedures and immediately transferred to the laboratory in holding medium composed of DMEM+GlutaMAX (Gibco) supplemented with 1% (v/v) PenStrep (Anprotec), 1% (v/v) L-Glutamine (Gibco), and 2,5µg/ml Amphotericin (CHEPLA PHARM). All subsequent handling was performed promptly under sterile conditions in a laminar flow cabinet. To remove residual blood, skin samples were washed three times with Hank’s Balanced Salt Solution (HBSS, Gibco) containing 4% (v/v) antibiotic-antimycotic solution, followed by a final wash in PBS (Anprotec). The tissue was then cut into small squares (approximately 8 × 8 mm²) and transferred into a pre-prepared 12-well plate (see **Figure S23 A**). Each well contained two sterile absorbent pads overlaid with a sterile nylon membrane pre-soaked in skin culture medium (DMEM+GlutaMAX (Gibco), supplemented with 1% (v/v) PenStrep (Anprotec), 1% (v/v) L-Glutamine (Gibco), 2.5 µg/ml Amphotericin (CHEPLA PHARM), 100 U/ml insulin (Merck), and 1µM dexamethasone (Sigma). One skin piece was placed epidermis side up on each membrane. Each well was filled with skin medium to enable air-liquid interface culture. Skin samples were incubated at 37 °C in a humidified atmosphere with 5% CO₂ for up to 14 days. The culture medium was refreshed every 2–3 days.

### LNP Injection into *Ex Vivo* Skin

For LNP administration, skin pieces were placed epidermis-side up on a sterile filter (Figure 6 **A**). A total volume of 100 µl of LNP formulations, diluted at the indicated concentrations in PBS, was carefully injected at the dermal–hypodermal interface. After injection, the skin was carefully washed in PBS before being placed back onto the nylon membrane in the culture dish. Sham injections with 100 µl PBS served as treatment controls, and untreated skin served as an additional control. Following injection, all skin samples were cultured at 37°C in a humidified incubator with 5% CO₂ for the specified time points. Culture medium was replaced every 2–3 days.

### Sample analysis

#### Assessment of Cell Viability

Cell viability of macrophages and fibroblasts was assessed using the Cell Proliferation Kit II (XTT; Roche Diagnostics), following the manufacturer’s protocol. The assay is based on the metabolic activity of viable cells, which reduce the XTT reagent to a water-soluble formazan dye. The resulting colour intensity, measured spectrophotometrically, correlates directly with the number of metabolically active cells. Viability was expressed as the ratio of the change in optical density (ΔOD) relative to untreated control cells cultured without LNP exposure.

#### Real-Time Monitoring of Fibroblast Proliferation

Live-cell proliferation of fibroblasts was monitored using the Incucyte S3 live-cell imaging system (Sartorius & Essen, BioScience). Cells were seeded in 24-well plates and stimulated with LNPs as described in the *LNP Stimulation of Macrophages and Fibroblasts* section. Proliferation was tracked over a 72-hour period, with 12 images captured per well at 6-hour intervals. Cell confluence was quantified using Incucyte analysis software (Sartorius), which calculated changes in confluence over time based on the acquired images for each condition.

#### RNA Isolation and Quantitative Real-Time PCR

For cell culture experiments, total RNA was isolated directly from adherent macrophages and fibroblasts using the RNeasy™ Mini Kit (Qiagen), following the manufacturer’s instructions. In skin culture experiments, the subcutaneous fat layer was first carefully separated from the skin, and the dermis and epidermis were subsequently separated from the remaining skin tissue by digestion with dispase for 2 hours. Isolated fat and dermal tissues were homogenised, and total RNA was extracted using the ReliaPrep™ RNA Tissue Miniprep System (Promega), according to the manufacturer’s protocol. RNA concentration and purity were assessed using a NanoDrop ND-1000 spectrophotometer (NanoDrop Technologies, Wilmington, DE, USA). For cDNA synthesis, 250 ng of total RNA was reverse transcribed using the LunaScript™ RT SuperMix Kit (New England Biolabs), following the manufacturer’s guidelines. Quantitative real-time PCR was performed using Luna® Universal qPCR Master Mix (New England Biolabs) with gene-specific primers (listed in Table S7). All PCR products were designed to span intron-exon boundaries.

Gene expression levels were quantified using standard curves generated from cloned cDNA and normalised to RPS26 (for cell samples) or GAPDH (for skin-derived dermis and fat).

#### Cytokine quantification

Concentrations of human TNF, IL-1β, MCP-1, and IL-10 in cell-free supernatants from macrophage cultures were measured using corresponding ELISA Kits (Thermo Fisher Scientific), according to the manufacturer’s protocols.

#### Histology and lipid and adipocyte staining of skin samples

Skin samples were cryo-frozen (Tissue Freezing Medium^TM^, Leica) and paraffin-embedded and skin sections prepared for immunohistochemistry or immunofluorescence staining.

For morphometric analysis, paraffin-embedded skin tissue sections (5 µm) were deparaffinized, rehydrated, and stained with hematoxylin and eosin using standard protocols. Stained sections were dehydrated, mounted with coverslips, and imaged by brightfield microscopy Oil Red O and LipidTOX™ staining were performed on cryosections of frozen skin tissue (50 µm thick). Sections were fixed in 4% paraformaldehyde, rinsed with distilled water, and incubated either with freshly prepared Oil Red O working solution (0.3% Oil Red O in 60% isopropanol, Sigma-Aldrich) for 20 minutes or with HCS LipidTOX™ Neutral Lipid Stain (Invitrogen), diluted 1:1000 in PBS, for 30 minutes at room temperature. Following staining, sections were rinsed with PBS. For LipidTOX-stained samples, nuclei were counterstained with DAPI (1:500 in PBS) for 10 minutes at room temperature. After a final wash, slides were mounted and imaged using brightfield microscopy (Oil Red O) or fluorescence microscopy (LipidTOX).

Perilipin (PLIN-1) staining was performed on paraffin-embedded skin tissue sections (10 µm thick). Following deparaffinization, antigen retrieval was carried out in citrate buffer (pH 6.0) for 30 minutes. After blocking and washing, sections were incubated overnight at 4 °C with anti-PLIN-1 primary antibody (Abcam, 1:500 dilution). The next day, sections were incubated with Alexa Fluor 546–conjugated secondary antibody (Invitrogen, 1:500), followed by nuclear counterstaining with DAPI (Invitrogen, 1:500).

Microscopy of all samples was performed with a Keyence BZ-X800 microscope (Biorevo) equipped with a fluorescence and image capturing system.

#### TUNEL assay

TUNEL staining was performed on paraffin-embedded tissue sections (5 µm) using the ApopTag® Fluorescein *In Situ* Apoptosis Detection Kit (Merck), following the manufacturer’s protocol. Sections were deparaffinized, rehydrated, and processed according to kit instructions. Nuclei were counterstained with DAPI (Invitrogen, 1:500), and fluorescence imaging was carried out using a Keyence BZ-X800 microscope (Biorevo).

#### Quantification of LNP Penetration Through Skin

To assess the penetration of LNPs through skin, samples injected with fluorescein-loaded LNP1-FL were placed in transwell inserts (translucent, 0.4 µm pore size; Greiner Bio-One) within a 12-well plate. Both the lower chamber and the transwell insert were filled with a fixed volume of skin culture medium, allowing for air-liquid interface conditions. At the indicated time points, medium from the lower chamber was collected and replaced with the same volume of fresh medium. Fluorescence in the collected medium was quantified using a fluorescence plate reader (SinergyHT, BioTek).

#### Statistical analysis

Comparisons involving more than two groups time were analysed using two-way ANOVA followed by Tukey’s multiple comparisons test. Statistical calculations were performed using GraphPad Prism, version 10. P-values ≤ 0.05 were considered statistically significant. Levels of significance are indicated as follows: *P < 0.05; **P < 0.01; ***P < 0.001; ****P < 0.0001.

## ASSOCIATED CONTENT

### Supporting Information

The following files are available free of charge.

Supporting Information (PDF)

Cryo-TEM 3D image (GIF)

## Author Contributions

The manuscript was written through the contributions of all authors. All authors have approved the final version of the manuscript. ‡These authors contributed equally.

## ACKNOWLEDGMENT

The authors acknowledge the 3D4D2 consortium for fruitful discussions. This includes J. Snoep, L.M. Birkholtz, B. Klumperman, L. Koekemoer, U. Freudenberg and E. Vassileva.

## ABBREVIATIONS

AF4: asymmetrical flow-field flow fractionation
MD: multidetector
MALS: multi angle light scattering
DLS: dynamic light scattering
SLD: scattering length density
SAXS: small angle X-ray scattering
WAXS: wide angle X-ray scattering
CCR2: CC chemokine receptor 2
CCL2: CC chemokine ligand 2
CCR5: CC chemokine receptor 5
TLC: thin layer chromatography.

## Notes

Any additional relevant notes should be placed here.

## REFERENCES

(1) Tenchov, R.; Bird, R.; Curtze, A. E., Zhou, Q. Lipid nanoparticles─From liposomes to mRNA vaccine delivery, a landscape of research diversity and advancement. ACS Nano. 2021, 15(*11*), 16982–17015.

(2) Hou, X.; Zaks, T.; Langer, R., Dong, Y. Lipid nanoparticles for mRNA delivery. Nat. Rev. Mater. 2021, 6(*12*), 1078–1094.

(3) Xu, L.; Wang, X.; Liu, Y.; Yang, G.; Falconer, R. J., Zhao, C.-X. Lipid nanoparticles for drug delivery. Adv. NanoBiomed. Res. 2022, 2(*2*), 2100109.

(4) Doktorovova, S.; Souto, E. B., Silva, A. M. Nanotoxicology applied to solid lipid nanoparticles and nanostructured lipid carriers – A systematic review of in vitro data. Eur. J. Pharm. Biopharm. 2014, 87(*1*), 1–18.

(5) Mitchell, M. J.; Billingsley, M. M.; Haley, R. M.; Wechsler, M. E.; Peppas, N. A., Langer, R. Engineering precision nanoparticles for drug delivery. Nat. Rev. Drug Discov. 2021, 20(*2*), 101–124.

(6) Haddadzadegan, S.; Dorkoosh, F., Bernkop-Schnürch, A. Oral delivery of therapeutic peptides and proteins: Technology landscape of lipid-based nanocarriers. Adv. Drug Deliv. Rev. 2022, 182, 114097.

(7) Viegas, C.; Patrício, A. B.; Prata, J. M.; Nadhman, A.; Chintamaneni, P. K., Fonte, P. Solid lipid nanoparticles vs. nanostructured lipid carriers: A comparative review. Pharmaceutics. 2023, 15(*6*).

(8) Gordillo-Galeano, A., Mora-Huertas, C. E. Solid lipid nanoparticles and nanostructured lipid carriers: A review emphasizing on particle structure and drug release. Eur. J. Pharm. Biopharm. 2018, 133, 285–308.

(9) Khosa, A.; Reddi, S., Saha, R. N. Nanostructured lipid carriers for site-specific drug delivery. Biomed. Pharmacother. 2018, 103, 598–613.

(10) Lacerda, S. P.; Cerize, N. N. P., Ré, M. I. Preparation and characterization of carnauba wax nanostructured lipid carriers containing benzophenone-3. Int. J. Cosmet. Sci. 2011, 33(*4*), 312–321.

(11) Madureira, A. R.; Campos, D. A.; Fonte, P.; Nunes, S.; Reis, F.; Gomes, A. M.; Sarmento, B., Pintado, M. M. Characterization of solid lipid nanoparticles produced with carnauba wax for rosmarinic acid oral delivery. RSC Adv. 2015, 5(*29*), 22665–22673.

(12) Spinozzi, F.; Moretti, P.; Perinelli, D. R.; Corucci, G.; Piergiovanni, P.; Amenitsch, H.; Sancini, G. A.; Franzese, G., Blasi, P. Small-angle X-ray scattering unveils the internal structure of lipid nanoparticles. J. Colloid Interface Sci. 2024, 662, 446–459.

(13) Ishkitiev, N.; Micheva, M.; Miteva, M.; Gaydarova, S.; Tzachev, C.; Lozanova, V.; Lozanov, V.; Cheshmedzhieva, D.; Kandinska, M.; Ilieva, S.; Gargallo, R.; Baluschev, S.; Stoynov, S.; Dyankova-Danovska, T.; Nedelcheva-Veleva, M.; Landfester, K.; Mihaylova, Z., Vasilev, A. Nanoconfined chlorine-substituted monomethine cyanine dye with a propionamide function based on the thiazole orange scaffold—Use of a fluorogenic probe for cell staining and nucleic acid visualization. Molecules. 2024, 29(*24*).

(14) Kaupbayeva, B.; Murata, H.; Matyjaszewski, K.; Russell, A. J.; Boye, S., Lederer, A. A comprehensive analysis in one run – in-depth conformation studies of protein–polymer chimeras by asymmetrical flow field-flow fractionation. Chem. Sci. 2021, 12(*41*), 13848– 13856.

(15) Boye, S.; Ennen, F.; Scharfenberg, L.; Appelhans, D.; Nilsson, L., Lederer, A. From 1D rods to 3D networks: A biohybrid topological diversity investigated by asymmetrical flow field-flow fractionation. Macromolecules. 2015, 48(*13*), 4607–4619.

(16) Engelke, J.; Boye, S.; Tuten, B. T.; Barner, L.; Barner-Kowollik, C., Lederer, A. Critical assessment of the application of multidetection SEC and AF4 for the separation of single-chain nanoparticles. ACS Macro Lett. 2020, 9(*11*), 1569–1575.

(17) Gumz, H.; Boye, S.; Iyisan, B.; Krönert, V.; Formanek, P.; Voit, B.; Lederer, A., Appelhans, D. Toward functional synthetic cells: In-depth study of nanoparticle and enzyme diffusion through a cross-linked polymersome membrane. Adv. Sci. 2019, 6(*7*), 1801299.

(18) Basson, I., Reynhardt, E. C. An investigation of the structures and molecular dynamics of natural waxes. II. Carnauba wax. J. Phys. D Appl. Phys. 1988, 21(*9*), 1429.

(19) Kawaguchi, T. Radii of gyration and scattering functions of a torus and its derivatives. J. Appl. Crystallogr. 2001, 34(*5*), 580–584.

(20) Gorzkiewicz, M.; Appelhans, D.; Boye, S.; Lederer, A.; Voit, B., Klajnert-Maculewicz, B. Effect of the structure of therapeutic adenosine analogues on stability and surface electrostatic potential of their complexes with poly(propyleneimine) dendrimers. Macromol. Rapid Commun. 2019, 40(*15*), 1900181.

(21) Boye, S.; Polikarpov, N.; Appelhans, D., Lederer, A. An alternative route to dye– polymer complexation study using asymmetrical flow field-flow fractionation. J. Chromatogr. A,. 2010, 1217(29), 4841–4849.

(22) Buechler, M. B.; Fu, W., Turley, S. J. Fibroblast-macrophage reciprocal interactions in health, fibrosis, and cancer. Immunity. 2021, 54(*5*), 903–915.

(23) Franklin, R. A. Fibroblasts and macrophages: Collaborators in tissue homeostasis. Immunol. Rev. 2021, 302(*1*), 86–103.

(24) Torregrossa, M.; Davies, L.; Hans-Günther, M.; Simon, J. C.; Franz, S., Rinkevich, Y. Effects of embryonic origin, tissue cues and pathological signals on fibroblast diversity in humans. Nat. Cell Biol. 2025, 27(*5*), 720–735.

(25) Franz, S.; Rammelt, S.; Scharnweber, D., Simon, J. C. Immune responses to implants – A review of the implications for the design of immunomodulatory biomaterials. Biomaterials. 2011, 32(*28*), 6692–6709.

(26) Wynn, T. A., Vannella, K. M. Macrophages in tissue repair, regeneration, and fibrosis. Immunity. 2016, 44(*3*), 450–462.

(27) Rodríguez-Morales, P., Franklin, R. A. Macrophage phenotypes and functions: resolving inflammation and restoring homeostasis. Trends Immunol. 2023, 44(*12*), 986–998.

(28) Atwood, S. X., Plikus, M. V. Fostering a healthy culture: Biological relevance of *in vitro* and *ex vivo* skin models. Exp. Dermatol. 2021, 30(*3*), 298–303.

(29) Neil, J. E.; Brown, M. B., Williams, A. C. Human skin explant model for the investigation of topical therapeutics. Sci. Rep. 2020, 10(*1*), 21192.

(30) Gou, S., Kalia, Y. N. Development of an *ex vivo* human skin model and evaluation of biological responses to subcutaneously injected hyaluronic acid formulations. Int. J .Pharm. 2025, 674, 125490.

(31) Grabner, G. F.; Xie, H.; Schweiger, M., Zechner, R. Lipolysis: cellular mechanisms for lipid mobilization from fat stores. Nat. Metab. 2021, 3(*11*), 1445–1465.

(32) Deng, J.; Liu, S.; Zou, L.; Xu, C.; Geng, B., Xu, G. Lipolysis response to endoplasmic reticulum stress in adipose cells. J Biol Chem. 2012, 287(*9*), 6240–6249.

(33) Zechner, R.; Zimmermann, R.; Eichmann, T. O.; Kohlwein, S. D.; Haemmerle, G.; Lass, A., Madeo, F. Fat signals - Lipases and lipolysis in lipid metabolism and signaling. Cell Metab. 2012, 15(*3*), 279–291.

(34) Brasaemle, D. L. Thematic review series: adipocyte biology. The perilipin family of structural lipid droplet proteins: stabilization of lipid droplets and control of lipolysis. J Lipid Res. 2007, 48(*12*), 2547–2559.

(35) Li, S.; Xue, T.; He, F.; Liu, Z.; Ouyang, S.; Cao, D., Wu, J. A time-resolved proteomic analysis of transcription factors regulating adipogenesis of human adipose derived stem cells. Biochem. Biophys. Res. Commun.. 2019, 511(*4*), 855–861.

(36) Ruan, H., Dong, L. Q. Adiponectin signaling and function in insulin target tissues. J. Mol. Cell. Biol. 2016, 8(*2*), 101–109.

(37) Schweiger, M.; Eichmann, T. O.; Taschler, U.; Zimmermann, R.; Zechner, R., Lass, A. Chapter Ten - Measurement of Lipolysis. In Methods in Enzymology; MacDougald OA, Ed.; Vol. 538; Academic Press, 2014; pp 171–193. DOI: 10.1016/B978-0-12-800280-3.00010-4.

(38) Dolatabadi, J. E. N.; Valizadeh, H., Hamishehkar, H. Solid Lipid Nanoparticles as Efficient Drug and Gene Delivery Systems: Recent Breakthroughs. Adv. Pharm. Bull. 2015, 5(*2*), 151–159.

(39) Verma, D. D.; Verma, S.; Blume, G., Fahr, A. Particle size of liposomes influences dermal delivery of substances into skin. Int. J .Pharm. 2003, 258(*1*), 141–151.

(40) Adib, Z. M.; Ghanbarzadeh, S.; Kouhsoltani, M.; Yari Khosroshahi, A., Hamishehkar, H. The Effect of Particle Size on the Deposition of Solid Lipid Nanoparticles in Different Skin Layers: A Histological Study. Adv. Pharm. Bull. 2016, 6(*1*), 31–36.

(41) SasView. SasView for Small Angle Scattering Analysis. [Available from: https://www.sasview.org/.]

(42) National Institute of Standards and Technology (NIST). NCNR Activation Calculator Tool.: NIST Center for Neutron Research; [Available from: https://www.ncnr.nist.gov/resources/activation/.]

(43) Lohmann, N.; Schirmer, L.; Atallah, P.; Wandel, E.; Ferrer, R. A.; Werner, C.; Simon, J. C.; Franz, S., Freudenberg, U. Glycosaminoglycan-based hydrogels capture inflammatory chemokines and rescue defective wound healing in mice. Sci. Transl. Med. 2017, 9(*386*), eaai9044.

(44) Franz, S.; Allenstein, F.; Kajahn, J.; Forstreuter, I.; Hintze, V.; Möller, S., Simon, J. C. Artificial extracellular matrices composed of collagen I and high-sulfated hyaluronan promote phenotypic and functional modulation of human pro-inflammatory M1 macrophages. Acta Biomater. 2013, 9(*3*), 5621–5629.

(45) Wilkinson, H. N.; Kidd, A. S.; Roberts, E. R., Hardman, M. J. Human Ex vivo Wound Model and Whole-Mount Staining Approach to Accurately Evaluate Skin Repair. JoVE - J. Vis. Exp. 2021, (168), e62326.

