## Supporting Information for "Disc-Toroid Hybrid Lipid Nanoparticles for Efficient Drug Encapsulation and Subcutaneous Delivery"

**Size and stability characterisation of LNPs using batch-DLS**

**Table S1:** Size and polydispersity index (PDI) of the various LNP formulations as determined by DLS over time

| Sample | Aqueous media | Time (months) | Cumulants Method | | PSD analysis (Intensity) |
|---|---|---|---|---|---|
| | | | z-average (d.nm)[a] | PDI[a] | Size (d.nm)[a, b] |
| $LNP_5$ | 0.9% NaCl | $t_0$ | 39.4 ± 0.2 | 0.171 ± 0.007 | 43.5 ± 1.7 |
| | | $t_5$ | 36.2 ± 0.1 | 0.090 ± 0.010 | 39.9 ± 0.4 |
| $LNP_5$-WS | 0.9% NaCl | $t_0$ | 35.8 ± 0.5 | 0.085 ± 0.014 | 39.2 ± 0.3 |
| $LNP_5$-Q | 0.9% NaCl | $t_0$ | 35.3 ± 0.0 | 0.065 ± 0.003 | 37.9 ± 0.0 |
| | | $t_5$ | 35.3 ± 0.1 | 0.058 ± 0.007 | 37.9 ± 0.2 |
| $LNP_5$-DHA | 0.9% NaCl | $t_0$ | 35.3 ± 0.1 | 0.087 ± 0.021 | 38.9 ± 1.0 |
| | | $t_5$ | 35.8 ± 0.1 | 0.103 ± 0.010 | 39.8 ± 0.3 |
| $LNP_5$-B3 | 0.9% NaCl | $t_0$ | 35.7 ± 0.2 | 0.089 ± 0.010 | 39.1 ± 0.1 |
| | | $t_{18}$ | 34.8 ± 0.2 | 0.038 ± 0.016 | 36.7 ± 0.2 |
| $LNP_1$ | 0.9% NaCl | $t_0$ | 37.6 ± 0.2 | 0.116 ± 0.005 | 42.2 ± 0.3 |
| | | $t_{18}$ | 38.1 ± 0.1 | 0.126 ± 0.006 | 42.2 ± 0.8 |
| $LNP_1$-Q | 0.9% NaCl | $t_0$ | 33.9 ± 0.1 | 0.082 ± 0.007 | 37.1 ± 0.4 |
| | | $t_{18}$ | 35.7 ± 0.3 | 0.151 ± 0.017 | 38.9 ± 0.5 |
| $LNP_1$-W | ultrapure water | $t_0$ | 32.8 ± 0.1 | 0.081 ± 0.004 | 35.9 ± 0.3 |
| | | $t_{18}$ | 33.1 ± 0.2 | 0.148 ± 0.012 | 37.4 ± 1.0 |
| $LNP_1$-CW-TPGS | ultrapure water | $t_0$ | 35.0 ± 0.1 | 0.145 ± 0.012 | 40.2 ± 0.6 |
| | | $t_{18}$ | 35.4 ± 0.4 | 0.175 ± 0.011 | 40.3 ± 1.0 |
| $LNP_1$-CW-T40 | ultrapure water | $t_0$ | 48.2 ± 0.2 | 0.167 ± 0.006 | 58.5 ± 0.8 |
| | | $t_{18}$ | 48.5 ± 0.1 | 0.164 ± 0.006 | 58.8 ± 0.5 |
| $LNP_1$-TPGS | ultrapure water | $t_0$ | 38.4 ± 0.1 | 0.193 ± 0.008 | 46.0 ± 1.2 |
| | | $t_{18}$ | 39.6 ± 0.2 | 0.197 ± 0.005 | 45.9 ± 1.7 |
| $LNP_1$-T40 | ultrapure water | $t_0$ | 39.3 ± 0.1 | 0.133 ± 0.006 | 45.8 ± 0.4 |
| | | $t_{18}$ | 40.3 ± 0.2 | 0.128 ± 0.013 | 46.2 ± 0.8 |
| $LNP_1$-Fl | 0.9% NaCl | - | 39.4 ± 0.3 | 0.176 ± 0.005 | 43.1 ± 1.1 |

*(a) The average and mean standard deviation of the z-average, polydispersity index (PDI), and particle size (d.nm ) are calculated from 3-5 replicates. b) The particle size (d.nm) is intensity-based. The hydrodynamic diameter is calculated via the Stokes-Einstein equation, which is only applicable to spherical particles.*

**Table S2:** Batch-to-batch comparison of LNP formulation by DLS

| Sample | Aqueous media | Cumulants Method | | PSD analysis (Intensity) |
|---|---|---|---|---|
| | | z-average (d.nm)[a] | PDI[a] | Size (d.nm)[a,b] |
| $LNP_5$ | 0.9% NaCl | 39.4 ± 0.2 | 0.171 ± 0.007 | 43.5 ± 1.7 |
| $LNP_5$-B2 | 0.9% NaCl | 36.4 ± 0.2 | 0.083 ± 0.010 | 39.7 ± 0.3 |
| $LNP_5$-B3 | 0.9% NaCl | 35.7 ± 0.2 | 0.089 ± 0.010 | 39.1 ± 0.1 |
| $LNP_5$-B4 | 0.9% NaCl | 35.6 ± 0.2 | 0.085 ± 0.007 | 38.9 ± 0.4 |
| $LNP_5$-Q | 0.9% NaCl | 35.3 ± 0.0 | 0.065 ± 0.003 | 37.9 ± 0.0 |
| $LNP_5$-Q-B2 | 0.9% NaCl | 32.9 ± 0.2 | 0.044 ± 0.017 | 34.9 ± 0.3 |
| $LNP_5$-Q-B4 | 0.9% NaCl | 34.2 ± 0.1 | 0.084 ± 0.009 | 37.4 ± 0.3 |
| $LNP_5$-DHA | 0.9% NaCl | 35.3 ± 0.1 | 0.087 ± 0.021 | 38.9 ± 1.0 |
| $LNP_5$-DHA-B2 | 0.9% NaCl | 34.0 ± 0.1 | 0.047 ± 0.009 | 36.2 ± 0.2 |
| $LNP_5$-DHA-B4 | 0.9% NaCl | 35.3 ± 0.1 | 0.062 ± 0.014 | 37.9 ± 0.4 |

*(a) The average and mean standard deviation of the z-average, polydispersity index (PDI), and particle size (d.nm ) are calculated from 3-5 replicates. (b) The particle size (d.nm) is intensity-based.*

**Table S3:** Zeta (ζ) potential values for the various LNP formulations

| Sample | Aqueous media | Zeta potential (mV)[a] | Conductivity (mS/cm) |
|---|---|---|---|
| $LNP_5$ | 1.5mM NaCl | -8.90 ± 0.57 | 0.146 ± 0.001 |
| $LNP_5$-Q | 1.5 mM NaCl | -15.10 ± 0.75 | 0.157 ± 0.001 |
| $LNP_5$-DHA | 1.5mM NaCl | -9.27 ± 0.37 | 0.150 ± 0.001 |
| $LNP_1$ | 1.5 mM NaCl | -7.41 ± 1.59 | 0.229 ± 0.001 |
| $LNP_1$-Q | 1.5 mM NaCl | -12.50 ± 0.94 | 0.215 ± 0.000 |
| $LNP_1$-W [b] | 1 mM NaCl | -8.41 ± 0.54 | 0.141 ± 0.001 |
| $LNP_1$-CW-TPGS | 1 mM NaCl | -10.90 ± 0.61 | 0.150 ± 0.001 |
| $LNP_1$-CW-T40 | 1 mM NaCl | -13.80 ± 0.35 | 0.151 ± 0.000 |
| $LNP_1$-TPGS | 1 mM NaCl | -11.50 ± 1.30 | 0.150 ± 0.000 |
| $LNP_1$-T40 | 1 mM NaCl | -11.70 ± 0.55 | 0.151 ± 0.000 |
| $LNP_1$-Fl | 1 mM NaCl | -11.40 ± 1.44 | 0.217 ± 0.001 |

*(a) Zeta potential measurements were performed at 25°C, with a monomodal analysis function, Samples $LNP_5$-WS with an auto-mode analysis function.*

**Table S4:** Size and polydispersity index of different sample concentrations of $LNP_1$-CW-TPGS as determined by batch-mode DLS for SAXS measurements

| Sample | Concentration[a] (mg.ml$^{-1}$) | Cumulants Method | | PSD analysis (Intensity) |
|---|---|---|---|---|
| | | z-average (d.nm)[b] | PDI[b] | Size (d.nm)[b,c] |
| $LNP_1$-CW-TPGS [d] | 11.02 | 29.9 ± 0.0 | 0.166 ± 0.006 | 34.4 ± 0.7 |
| | 4.80 | 31.5 ± 0.1 | 0.137 ± 0.005 | 36.5 ± 0.5 |
| | 2.42 | 31.6 ± 0.2 | 0.149 ± 0.010 | 36.6 ± 0.9 |
| | 0.48 | 35.4 ± 0.2 | 0.146 ± 0.011 | 40.0 ± 1.1 |

*(a) Concentration based on lipid fraction. (b) The average and mean standard deviation of the z-average, polydispersity index (PDI), and particle size (d.nm ) are calculated from 3-5 replicates. (c) The particle size (d.nm) is intensity-based. d) Aqueous media: MilliQ ultrapure water.*

### Characterisation of LNP using AF4-MD

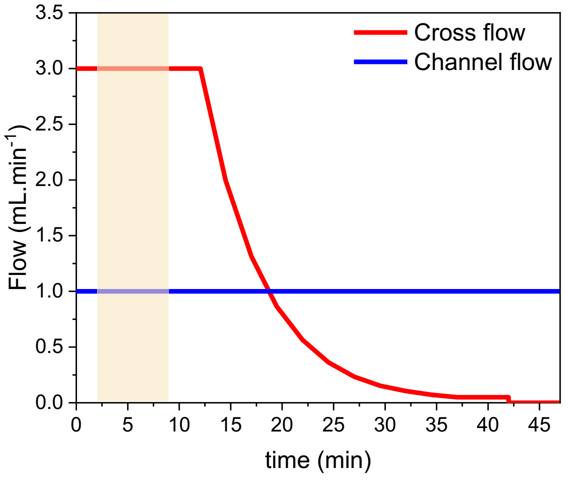


**Figure S1**. AF4 separation flow profile for LNP using 10 mM PBS at pH 7.4 as eluent.

**Table S5**: Experimental condition for the characterisation of LNP by AF4-MD

| **Conditions** | |
|---|---|
| Injection volume: | 25 µL and 250 µL (QELS) |
| Spacer: | 350 µm |
| Membrane: | 10 kDa (regenerate cellulose, Wyatt) |
| Detector flow: | 0.5 $mL.min^{-1}$ |
| Eluent: | 10 mM PBS buffer at pH 7.4, containing 200 $mg.L^{-1}$ $NaN_3$ |
| Channel: | Eclipse short channel with DCM, Wyatt Technologies Corp. |
| **Detector** | |
| Agilent 1260 Infinity II MWD | Wavelength: 250 nm, 280 nm, 300 nm, 310 nm, and 330 nm |
| Multi-angle light scattering (MALS) with online QELS (DAWN Neon MALLS, Wyatt Technologies Corp.) | 18 angles, operating at a wavelength of 659 nm |
| OptiLab dRI | Operating at a wavelength of 658 nm |
| Method: | Isocratic step with a $V_x$ of 3 $ml.min^{-1}$ for 9 min (Elution; Focus; Focus Inject; Elution).<br>Exponential $V_x$ gradient (slope 8) from 3 to 0.05 $mL.min^{-1}$ within 28 min was used.<br>Isocratic $V_x$ of 0.5 $mL.min^{-1}$ for 7 min.<br>Isocratic $V_x$ of 0 $mL.min^{-1}$ for 15 min.<br>Injection flow rate: 0.2 $mL.min^{-1}$ |

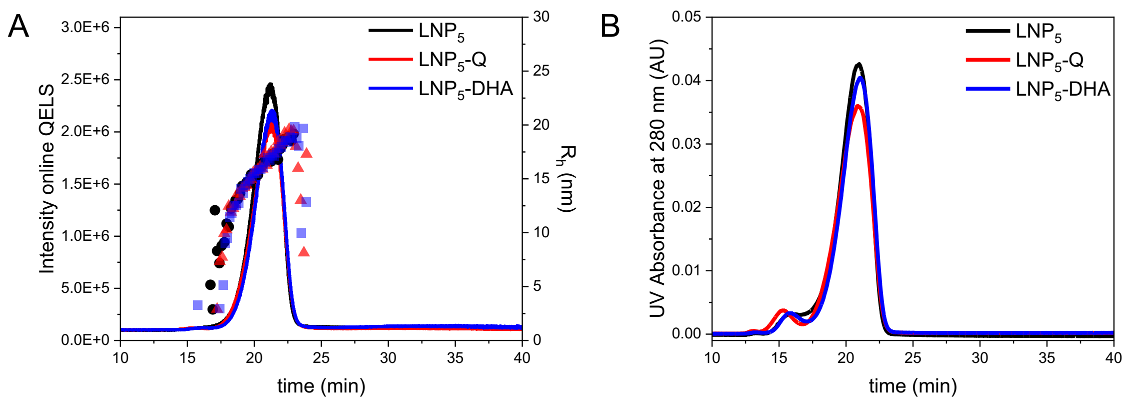


***Figure S2**. **(A)** Overlay of online QELS intensity vs. time for AF4 separation of $LNP_5$ unloaded, $LNP_5$-Q, and $LNP_5$-DHA, highlighting the $R_h$ distribution across the eluting peak. **(B)** Overlay of UV absorbance at 280 nm vs. time for AF4 separation of $LNP_5$ unloaded, $LNP_5$-Q, and $LNP_5$-DHA, illustrating the absorbance profiles among the formulations. Trace amounts of surfactant micelle visible at approx. 16 min*

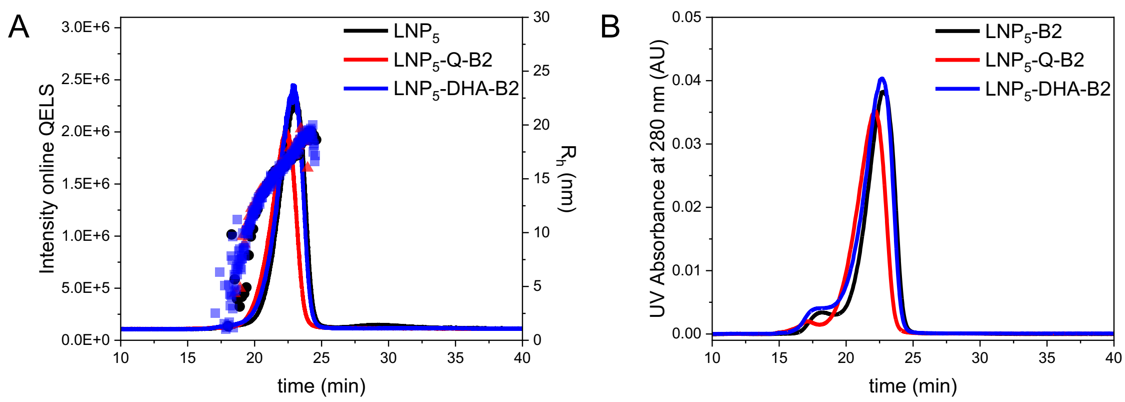


***Figure S3**. **(A)** Overlay of online QELS intensity vs. time for AF4 separation of $LNP_5$-B2 unloaded, $LNP_5$-Q-B2, and $LNP_5$-DHA-B2, highlighting the $R_h$ distribution across the eluting peak. **(B)** Overlay of UV absorbance at 280 nm vs. time for AF4 separation $LNP_5$ -B2 unloaded, $LNP_5$-Q-B2, and $LNP_5$-DHA-B2, illustrating the absorbance profiles among the formulations.*

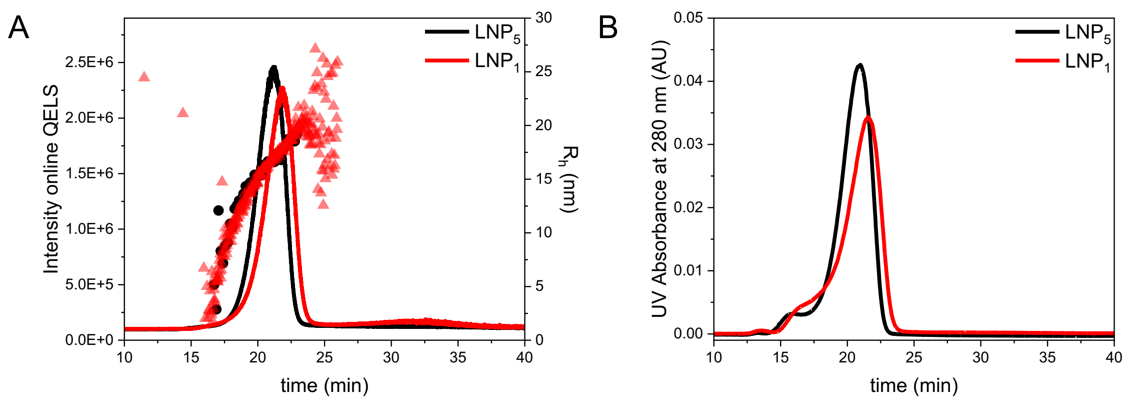


***Figure S4:** Illustrating the influence of the percentage lipid fraction on the size distribution **(A)** Overlay of online QELS intensity vs. time for AF4 separation of $LNP_5$ unloaded, 5.5 % lipid fraction and $LNP_1$ unloaded, 1.1 % lipid fraction. **(B)** Overlay of UV absorbance at 280 nm vs. time., trace amounts of surfactant micelle visible at approx. 16 min.*

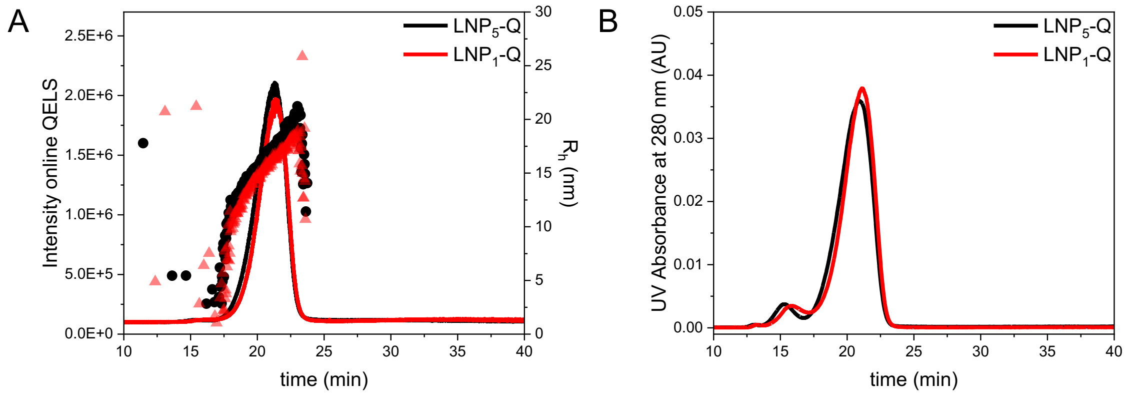


***Figure S5:*** *AF4 separation of $LNP_5$-Q, 5.5 % lipid fraction with 10% quinine incorporation and $LNP_1$–Q, 1.1 % lipid fraction with 10% quinine incorporation, to study the influence of the lipid fraction and percentage drug incorporation.* ***(A)*** *Overlay of online QELS intensity vs. time and* ***(B)*** *Overlay of UV absorbance at 280 nm vs. time, trace amounts of surfactant micelle visible at approx. 16 min .*

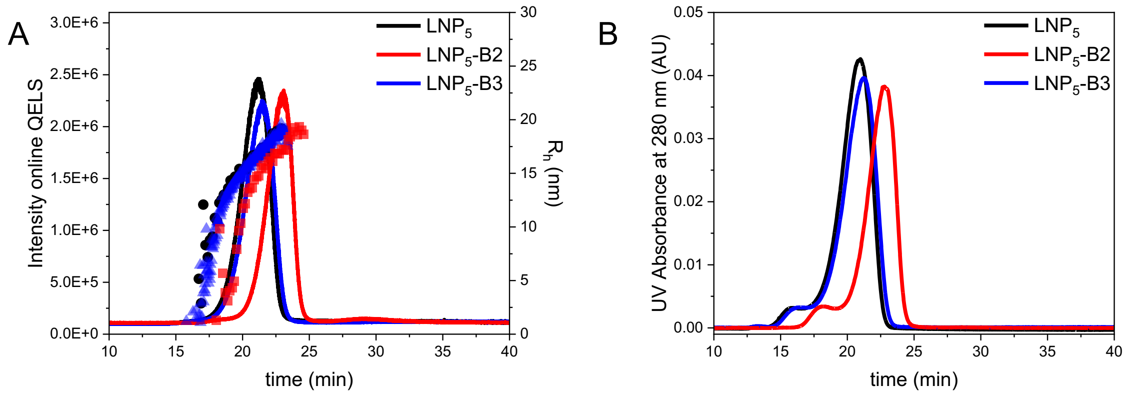


***Figure S6:*** *Batch-to-Batch comparison of the $LNP_5$ with 5.5% lipid fraction formulations.* ***(A)*** *Overlay of online QELS intensity vs. time for AF4 separation of $LNP_5$ for three different batch of LNP.* ***(B)*** *Overlay of UV absorbance at 280 nm vs. time*

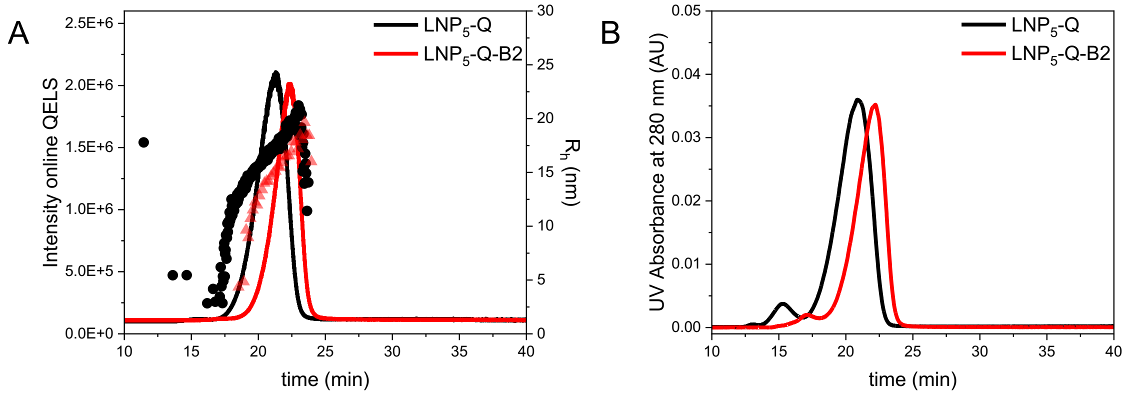


***Figure S7:*** *Batch-to-Batch comparison of the $LNP_5$ with quinine incorporation.* ***(A)*** *Overlay of online QELS intensity vs. time for AF4 separation of $LNP_5$ for two different batch of LNP.* ***(B)*** *Overlay of UV absorbance at 280 nm vs. time, trace amounts of surfactant micelle visible at approx. 15-17 min.*

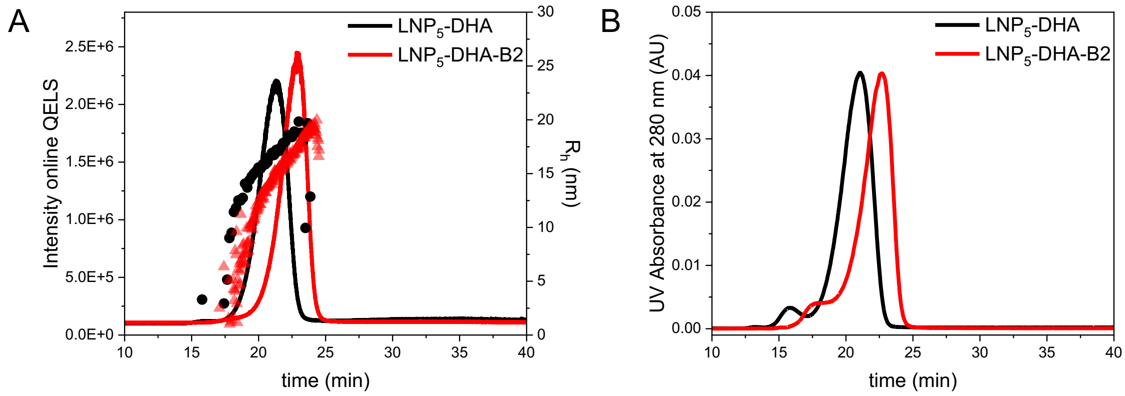


***Figure S8:*** *Batch-to-Batch comparison of the LNP$_5$ with dihydroartemisinin incorporation.* ***(A)*** *Overlay of online QELS intensity vs. time for AF4 separation of LNP$_5$-DHA for two different batch of LNP.* ***(B)*** *Overlay of UV absorbance at 280 nm vs. time, trace amounts of surfactant micelle visible at approx. 16 min..*

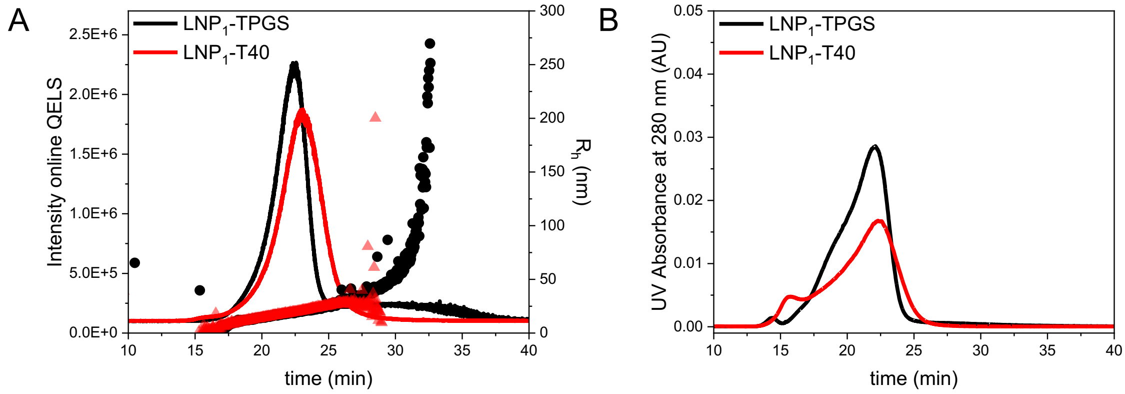


***Figure S9:*** *Comparison of AF4 separation of two LNP$_1$ samples with different surfactants, showing differences in size distribution over time.* ***(A)*** *Overlay of online QELS intensity vs. time for AF4 separation* ***(B)*** *Overlay of UV absorbance at 280 nm vs. time, trace amounts of surfactant micelle visible at approx. 16 min.*

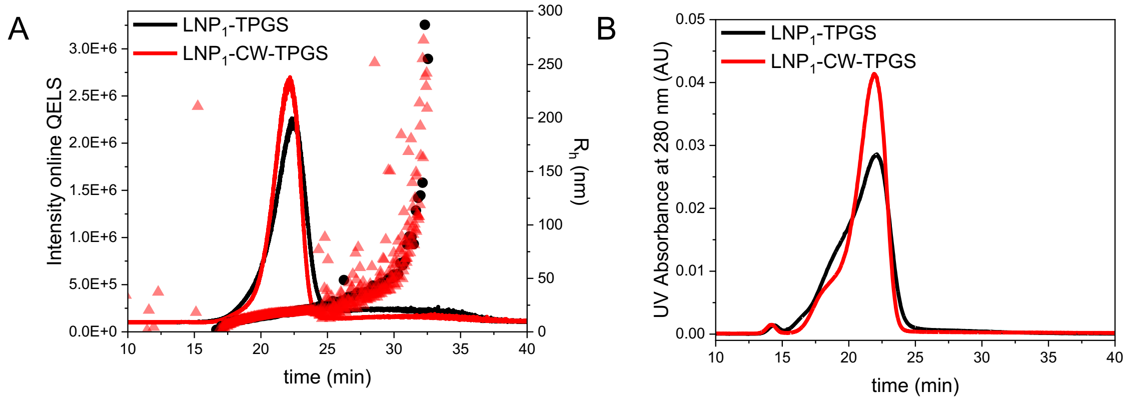


***Figure S10:*** *Comparison of AF4 separation of two LNP$_1$ samples with different lipid cores, but with the same surfactant (TPGS). LNP$_1$-TPGS has a lipid core that consist of carnauba wax and red palm oil, whereas LNP$_1$-CW-TPGS has a lipid core that consist of carnauba wax.* ***(A)*** *Overlay of online QELS intensity vs. time for AF4 separation showing differences in size distribution over time* ***(B)*** *Overlay of UV absorbance at 280 nm vs. time, trace amounts of surfactant micelle visible at approx. 16 min.*

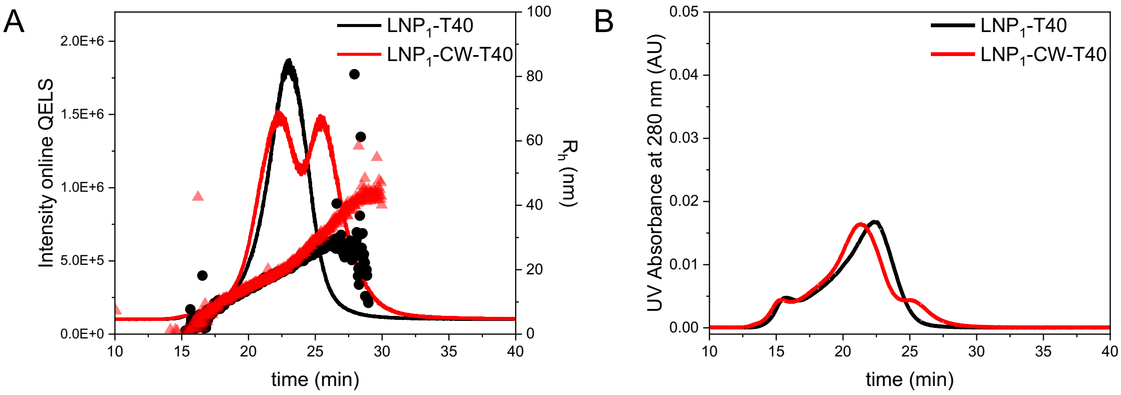


***Figure S11:*** *Comparison of AF4 separation of two LNP$_1$ samples with different lipid cores, but with similar surfactant (Polysorbate 40). LNP$_1$-T40 has a lipid core that consist of carnauba wax and red palm oil, whereas LNP$_1$-CW-T40 has a lipid core that consist of carnauba wax.* ***(A)*** *Overlay of online QELS intensity vs. time for AF4 separation showing differences in size distribution over time* ***(B)*** *Overlay of UV absorbance at 280 nm vs. time, trace amounts of surfactant micelle visible at approx. 16 min.*

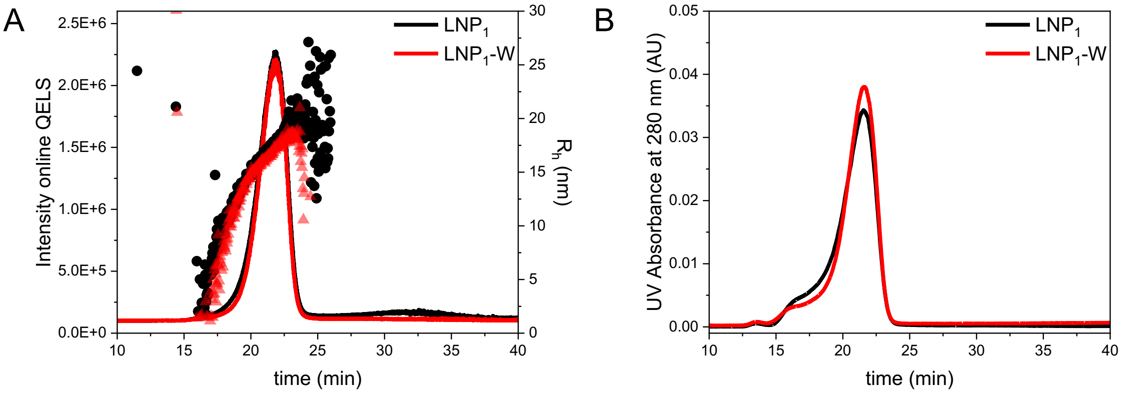


***Figure S12:*** *Comparison of AF4 separation of two LNP$_1$ formulations prepared in different dispersions. LNP$_1$ has been dispersed in 0.9% NaCl and LNP$_1$-W has been dispersed in ultrapure water.* ***(a)*** *Overlay of online QELS intensity vs. time for AF4 separation* ***(b)*** *Overlay of UV absorbance at 280 nm vs. time, trace amounts of surfactant micelle visible at approx. 16 min.*

**Table S6:** Summary of Radius of gyration ($R_g$), Hydrodynamic radius ($R_h$) and $R_g/R_h$ ratio for various LNP formulations and batches

| Sample | $R_g$ [(a)] | $R_h$ [(a)] | $\frac{R_g}{R_h}$ [(a)] |
|---|---|---|---|
| $LNP_5$ | 11.4 ± 0.3 | 16.8 ± 0.4 | 0.68 |
| $LNP_5$-Q | 10.4 ± 0.5 | 17.4 ± 0.4 | 0.60 |
| $LNP_5$-DHA | 10.6± 0.3 | 17.3 ± 0.4 | 0.61 |
| $LNP_1$ | 11.6 ± 0.3 | 17.9 ± 0.4 | 0.65 |
| $LNP_1$-Q | 10.8 ± 0.3 | 16.4 ± 0.4 | 0.66 |
| $LNP_1$-W | 11.2 ± 0.3 | 17.3 ± 0.4 | 0.64 |
| $LNP_1$-TPGS | 12.9 ± 0.3 | 18.8 ± 0.4 | 0.69 |
| $LNP_1$-T40 | 16.0 ± 0.2 | 20.5 ± 0.5 | 0.78 |
| $LNP_1$-CW-TPGS | 10.9 ± 0.3 | 17.5 ± 0.4 | 0.62 |
| $LNP_1$-CW-T40 (first eluting peak) | 14.7 ± 0.3 | 19.5 ± 0.4 | 0.75 |
| $LNP_1$-CW-T40 (second eluting peak) | 25.5 ± 0.2 | 30.5 ± 0.7 | 0.84 |
| $LNP_5$-B2 | 12.6 ± 0.4 | 17.3 ± 0.4 | 0.73 |
| $LNP_5$-Q-B2 | 12.2 ± 0.3 | 16.8 ± 0.4 | 0.73 |
| $LNP_5$-DHA-B2 | 12.4 ± 0.3 | 17.2 ± 0.4 | 0.72 |
| $LNP_5$-B3 | 10.8 ± 0.3 | 17.3 ± 0.4 | 0.62 |

[(a)] Determined at peak height

### Cryo-TEM characterisation of LNPs

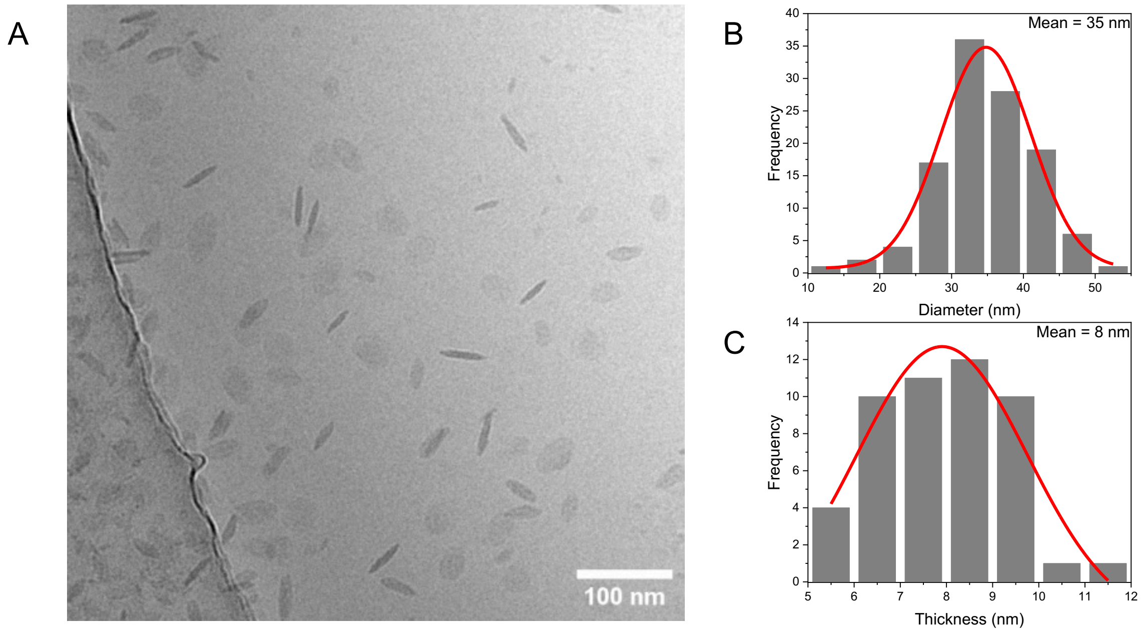


***Figure S13:*** *Cryo-TEM analysis of LNP5. **(A)** Cryo-TEM image of the particles. **(B)** Particle diameter distribution measured from 20-40 particles across three images, fitted with a Gaussian distribution. **(C)** Particle thickness distribution measured from 10-20 particles across three images, fitted with a Gaussian distribution.*

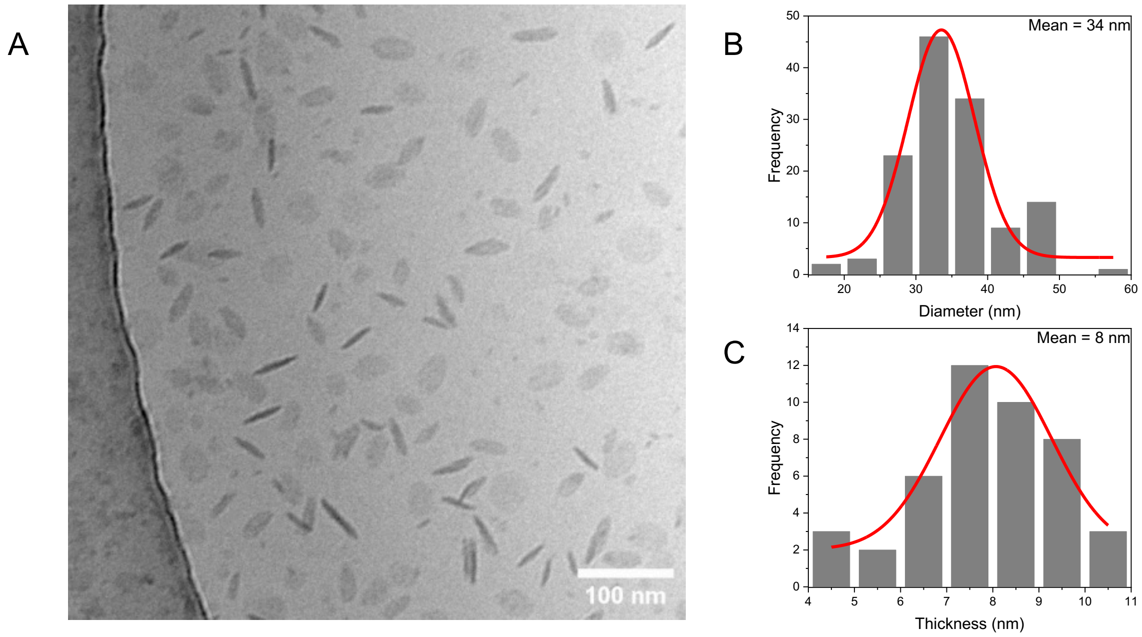


***Figure S14:*** *Cryo-TEM analysis of LNP5-Q. **(A)** Cryo-TEM image of the particles. **(B)** Particle diameter distribution measured from 20-40 particles across three images, fitted with a Gaussian distribution. **(C)** Particle thickness distribution measured from 10-20 particles across three images, fitted with a Gaussian distribution.*

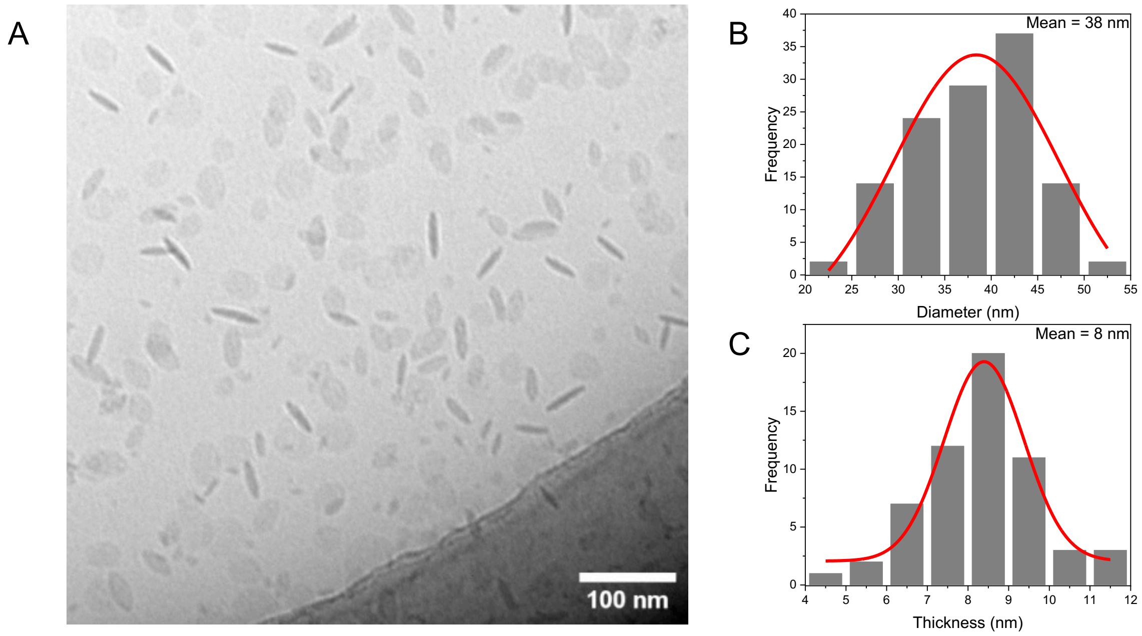


***Figure S15:*** *Cryo-TEM analysis of LNP$_5$-DHA.* ***(A)*** *Cryo-TEM image of the particles.* ***(B)*** *Particle diameter distribution measured from 20-40 particles across three images, fitted with a Gaussian distribution.* ***(C)*** *Particle thickness distribution measured from 10-20 particles across three images, fitted with a Gaussian distribution.*

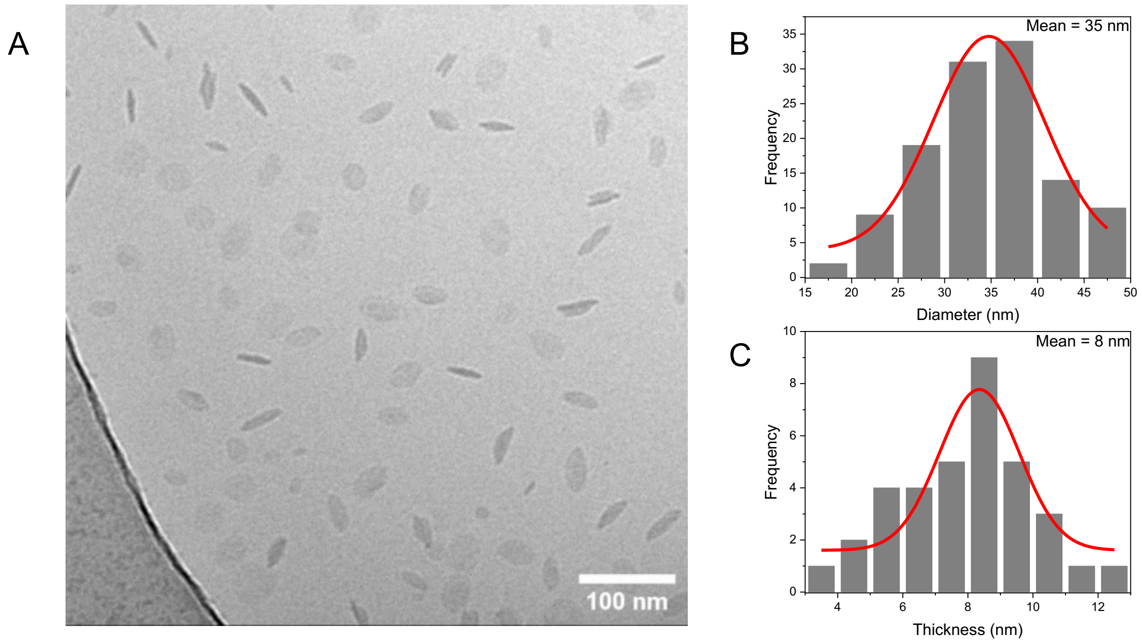


***Figure S16:*** *Cryo-TEM analysis of LNP$_1$-W.* ***(A)*** *Cryo-TEM image of the particles.* ***(B)*** *Particle diameter distribution measured from 20-40 particles across three images, fitted with a Gaussian distribution.* ***(C)*** *Particle thickness distribution measured from 10-20 particles across three images, fitted with a Gaussian distribution.*

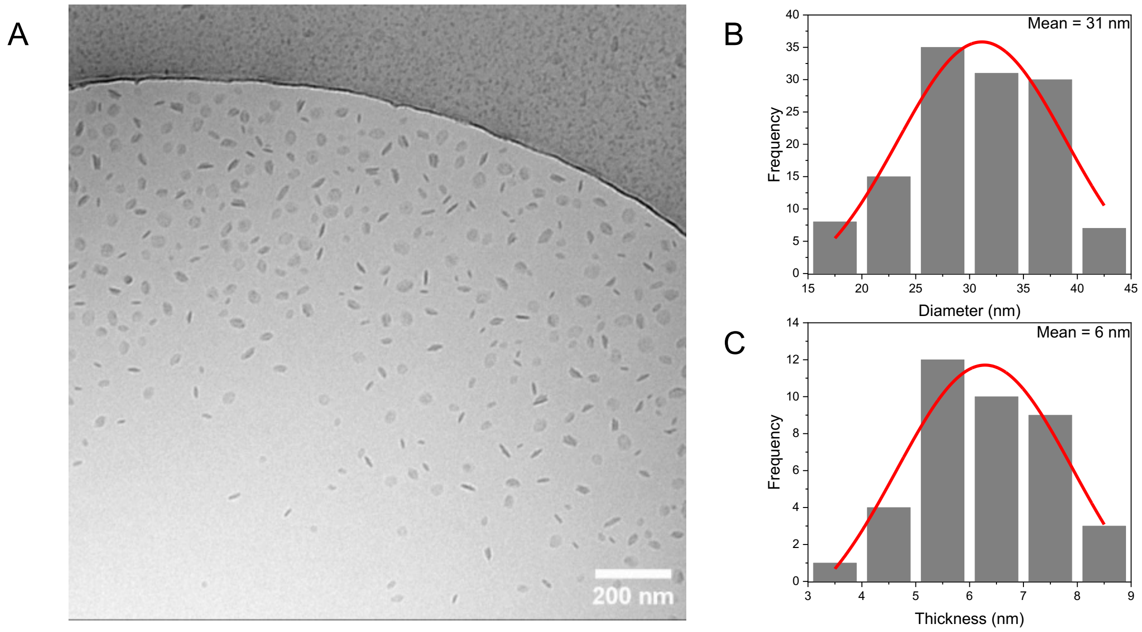


***Figure S17:*** *Cryo-TEM analysis of $LNP_1$-CW-TPGS.* ***(A)*** *Cryo-TEM image of the particles.* ***(B)*** *Particle diameter distribution measured from 20-40 particles across three images, fitted with a Gaussian distribution.* ***(C)*** *Particle thickness distribution measured from 10-20 particles across three images, fitted with a Gaussian distribution.*

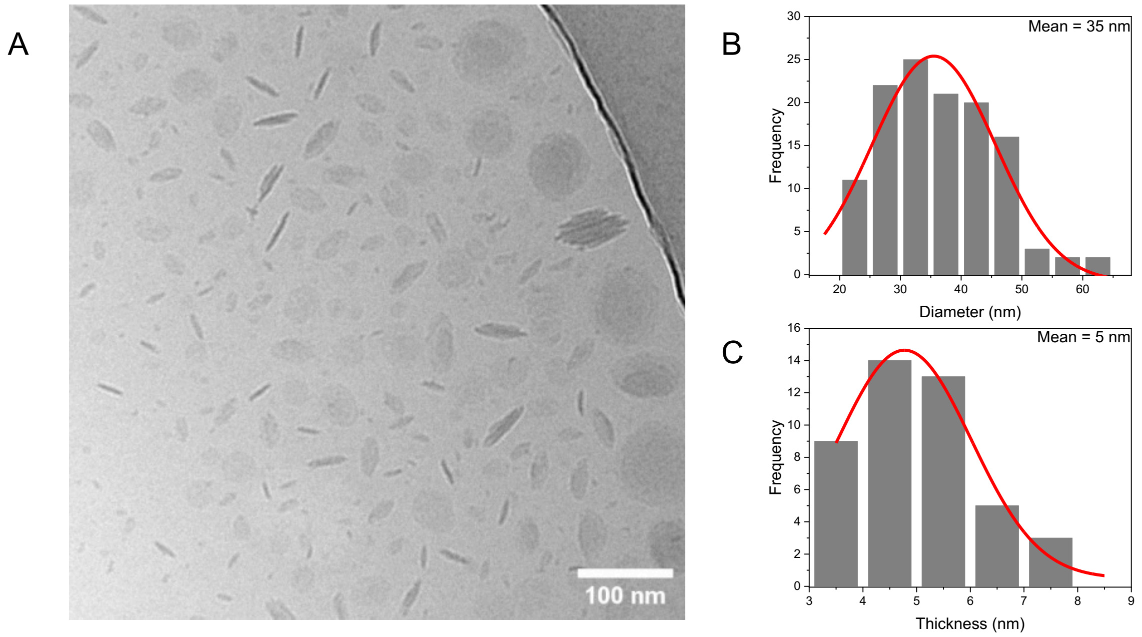


***Figure S18:*** *Cryo-TEM analysis of $LNP_1$-CW-T40.* ***(A)*** *Cryo-TEM image of the particles.* ***(B)*** *Particle diameter distribution measured from 20-40 particles across three images, fitted with a Gaussian distribution.* ***(C)*** *Particle thickness distribution measured from 10-20 particles across three images, fitted with a Gaussian distribution.*

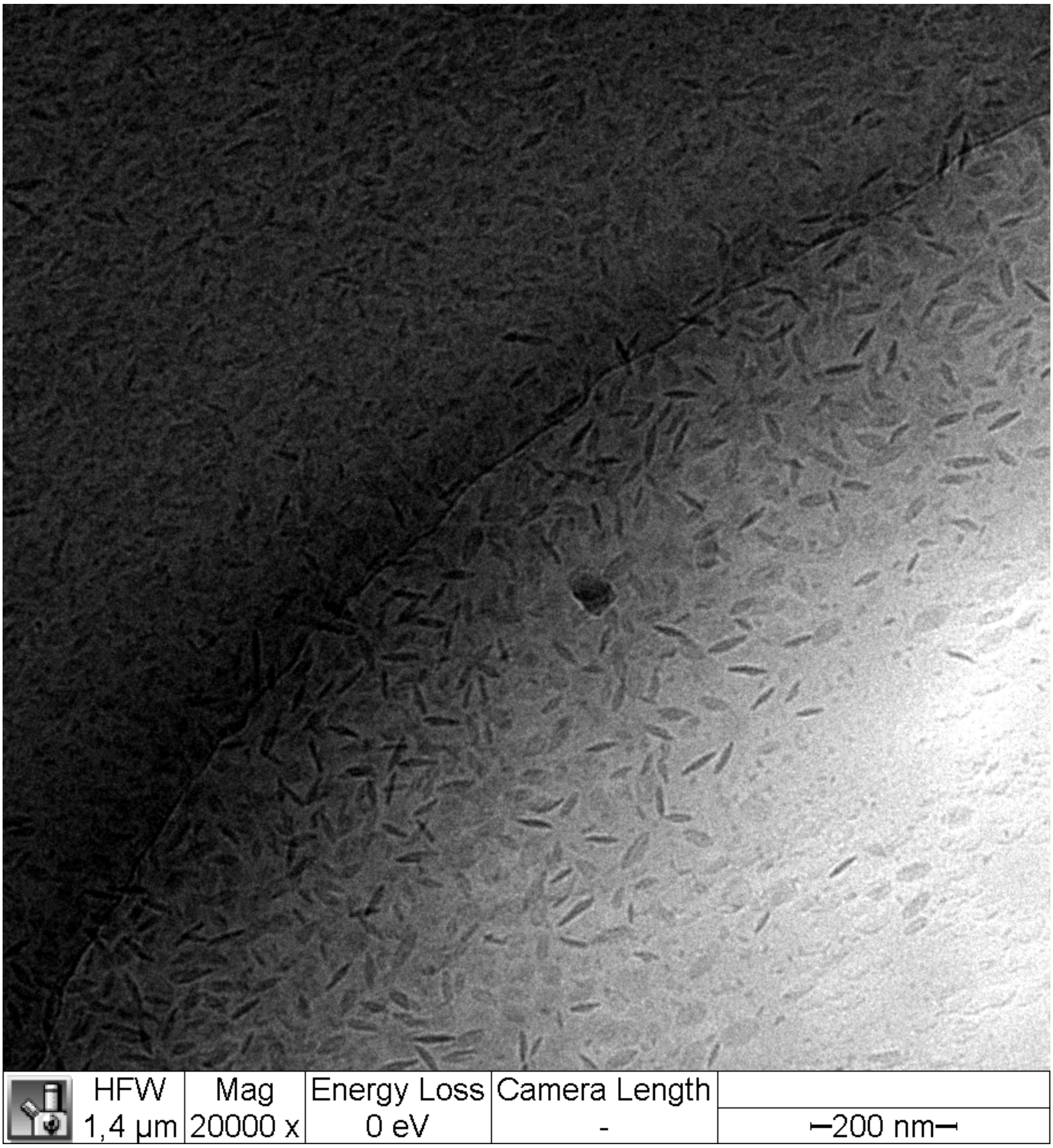


***Figure S20:*** *A three-dimensional rendering of the cryo-TEM micrograph of the LNPs. A video is provided as Supplementary Video 1.*

### Characterisation of LNP using SAXS

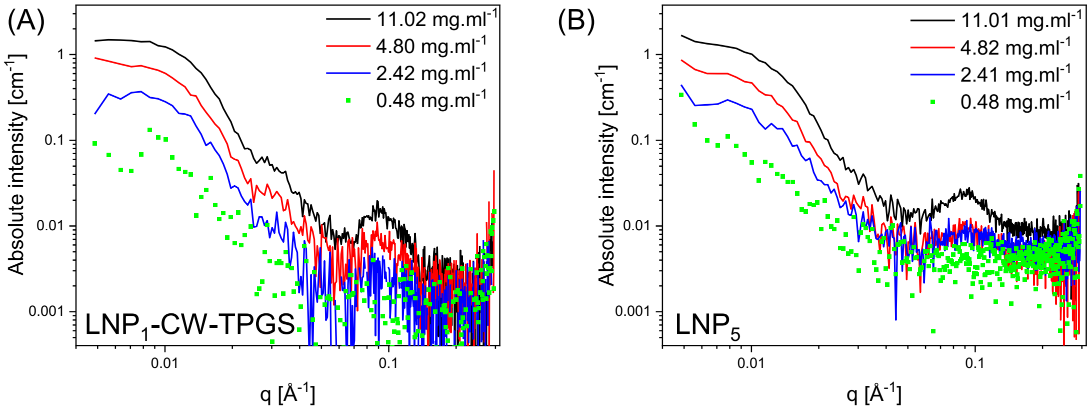


***Figure S21:*** *SAXS data converted to absolute intensity for LNP₁-CW-TPGS (A) and LNP₅ (B) to evaluate $R_g$ for four different concentrations of LNP₁-CW-TPGS and LNP₅ in ultrapure water.*

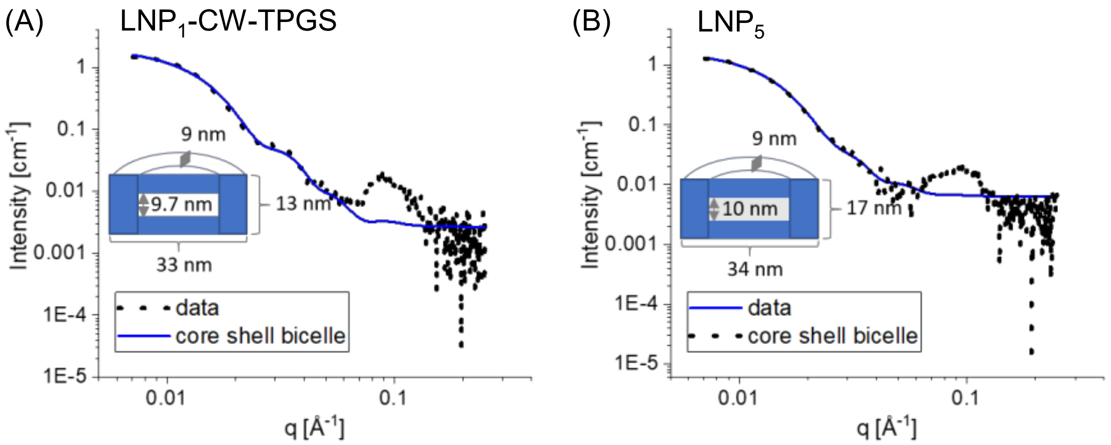


***Figure S22:*** *SAXS data and modelling by a core-shell bicelle for LNP₁-CW-TPGS (A) and LNP₅ (B) in water at a concentration of 11.02 mg.ml⁻¹ based on lipid fraction. The reflection at q = 0.09 Å⁻¹ cannot be captured by the model. Sketches of the modelled shape, considering two different SLDs of the core and the shell of the bicelle, are added to the graphs. For the LNP₁-CW-TPGS, which consists only of a carnauba wax core, the SLD of the core was close to that of the solvent water. In contrast, for the LNP₅, the SLD of the core was closer to that of carnauba wax.*

### Bioactivity and biocompatibility studies

**Table S7:** Human primer

| Gene | Accession number | Forward primer | Reverse primer |
|---|---|---|---|
| COL1a1 | NM_000088.4 | GTCGCACTGGTGATGCTG | GGTGGTGTCCACCTCGAG |
| COL3a1 | NM_000090.4 | CCATTGCTGGGATTGGAG | GTCCACCAGTGTTTCCGTG |
| EDA-FN | NM_002026.2 | CCAGTCCACAGCTATTCCTG | ACAACCACGGATGAGCTG |
| ACTA2(aSMA) | NM_001141945.3 | AGACCCTGTTCCAGCCATC | TGCTAGGGCCGTGATCTC |
| TNF | NM_000594.3 | GAGTGACAAGCCTGTAGCCCATGTTGTAGCA | GCAATGATCCCAAAGTAGACCTGCCCAGACT |
| IL-1b | NM_000576.3 | GACACATGGGATAACGAGGC | ACGCAGGACAGGTACAGATT |
| MCP1 | NM_002982.4 | CAG CCA GAT GCA ATC AAT GC | GTC TTC GGA GTT TGG GTT TGC |
| IL-10 | NM_000572.3 | GAT CTC CGA GAT GCC TTC AG | CAT GCG CCT TGA TGT CTG |
| PPARγ | NM_138712.5 | AGTCCTCACAGCTGTTTGCCAAGC | GAGCGGGTGAAGACTCATGTCTGTC |
| ADIPOQ | NM_001177800.2 | ACCCAGAGCTGTGGACTTTG | GTAGTCCTTCCAAGACAGGACG |
| PNPLA2 (ATGL) | NM_020376.4 | GAG TGA CAT CTG TCC GCA GG | GCG AGT AAT CCT CCG CTT GG |
| RPS26 | NM_001029 | CAATGGTCGTGCCAAAAAG | TTCACATACAGCTTGGGAAGC |
| GAPDH | NM_002046.7 | CTTCACCACCATGGAGAAGGC | CCAGTGAGCTTCCCGTTCAGC |

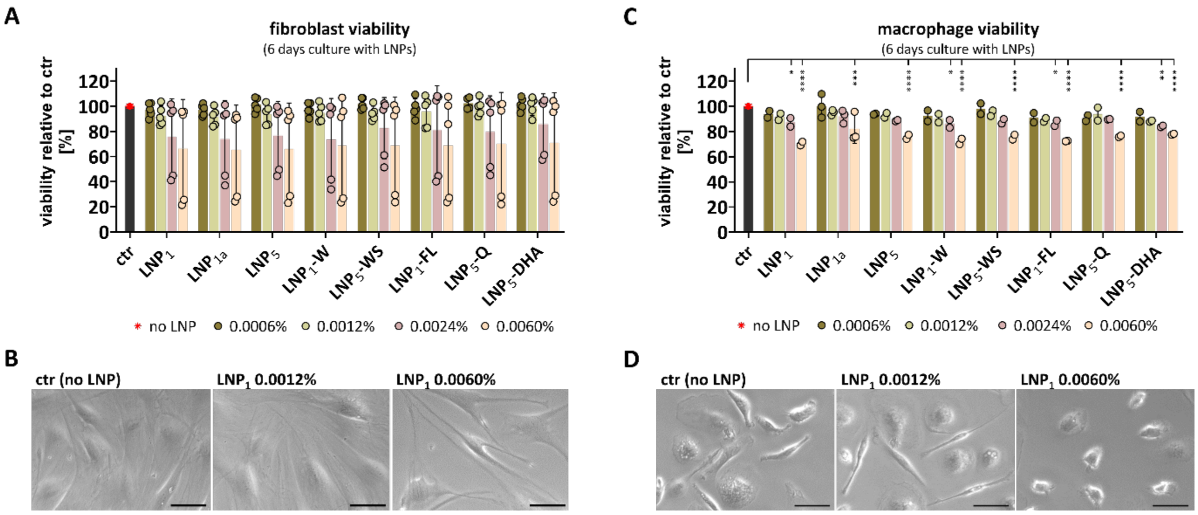


***Figure S21: Cell viability after culture with LNP.*** *Human dermal fibroblasts and monocyte-derived human macrophages were cultured for 6 days with different LNPs at different concentrations.* ***(A)*** *Viability of fibroblasts was determined with the XTT assay. n=4 different fibroblast donors.* ***(B)*** *Microscopic evaluation of fibroblasts. Scale = 50 µm.* ***(C)*** *Viability of macrophages was determined with the XTT assay. n=3 different macrophage donors.* ***(D)*** *Microscopic evaluation of macrophages. Scale = 50 µm. A/C) Two-way ANOVA with Tukey's multiple comparisons test. Significant differences compared to ctr (no LNP) are indicated. * p<0.05, *** p<0.001, **** p<0.0001. LNP concentrations are indicated as % lipid fraction. $LNP_1$, $LPN_{1a}$, $LNP_5$ comprise different batches of LNP constituted in 0.9% NaCl at stock concentrations of 1.1%, 1.2% and 5.5% lipid fraction, respectively. $LNP_1$-W are constituted in pure water. $LNP_5$-WS, the residual surfactants were removed. $LNP_1$-FL, $LNP_5$-Q, and $LNP_5$-DHA comprise batches of $LNP_1$ loaded with fluorescein (FL) for LNP tracking or with the drugs quinine (Q) or dihydroartemisinin (DHA).*

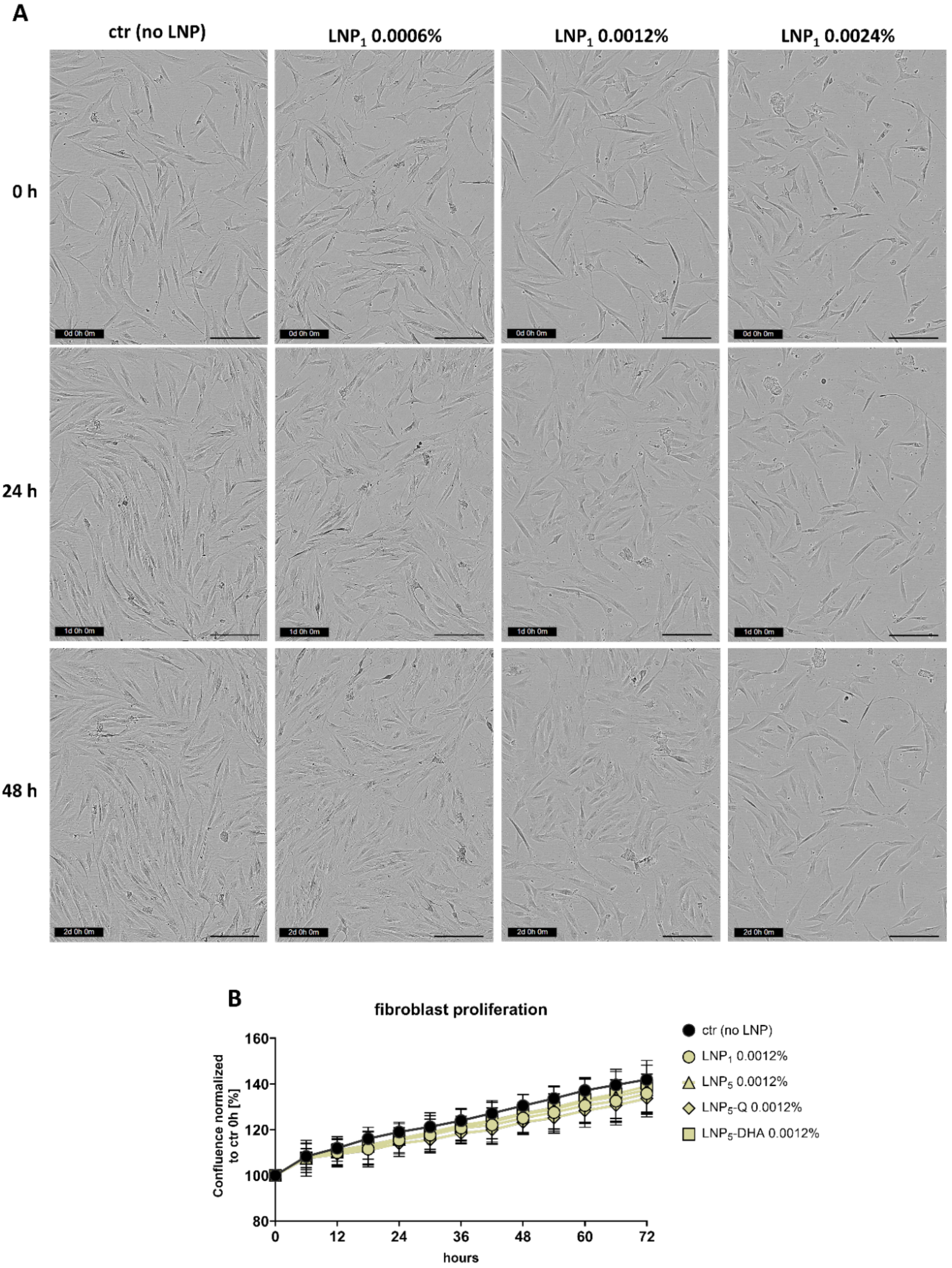


***Figure S22: Proliferation of fibroblasts monitored via IncuCyte® live cell imaging****. Human dermal fibroblasts were incubated with various LNP at different concentrations in the IncuCyte® S3. For a total of 72 h, 12 defined areas in each well with fibroblasts were photographed at a time interval of 6 h and used to determine fibroblast proliferation over 72 h.* ***(A)*** *Images of one representative area for the conditions indicated, photographed at 0 hours, 24 hours and 48 hours are shown (refers to* ***Figure 5A****). Scale = 200 µm.* ***(B)*** *Quantification of fibroblast proliferation with different LNP. n=4. Two-way ANOVA with Dunnett's multiple comparisons test was performed. No significant differences were observed. LNP concentrations are indicated as % lipid fraction. LNP$_1$, LNP$_5$, LNP$_5$-Q, LNP$_5$-DHA comprise different batches of LNP as outlined in* ***Figure 4****.*

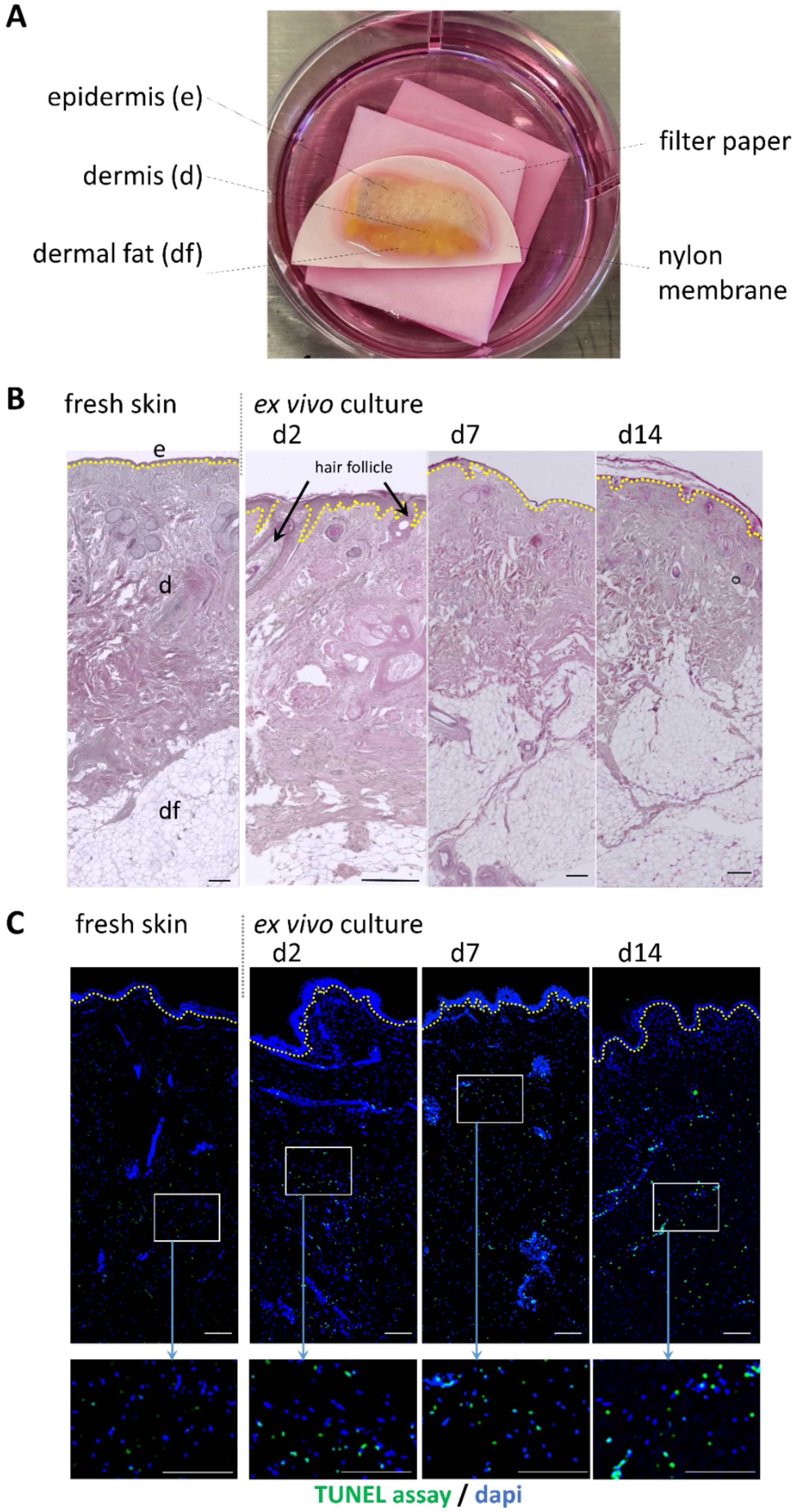


***Figure S23: Ex vivo skin culture model.*** ***(A)*** *Image of human skin prepared for ex vivo culture in a culture plate on two filter papers and a nylon membrane to ensure adequate medium and nutrient supply.* ***(B)*** *Hematoxylin and Eosin (H&E) staining of histology sections of fresh human skin and human skin cultured ex vivo for two days, 7 days and 14 days. scale = 300μm* ***(C)*** *Evaluation of skin viability via TUNEL staining of histology sections of fresh human skin and human skin cultured ex vivo for two days, 7 days and 14 days. scale = 200μm.*

### Characterisation of LNP loading using fluorescence spectroscopy

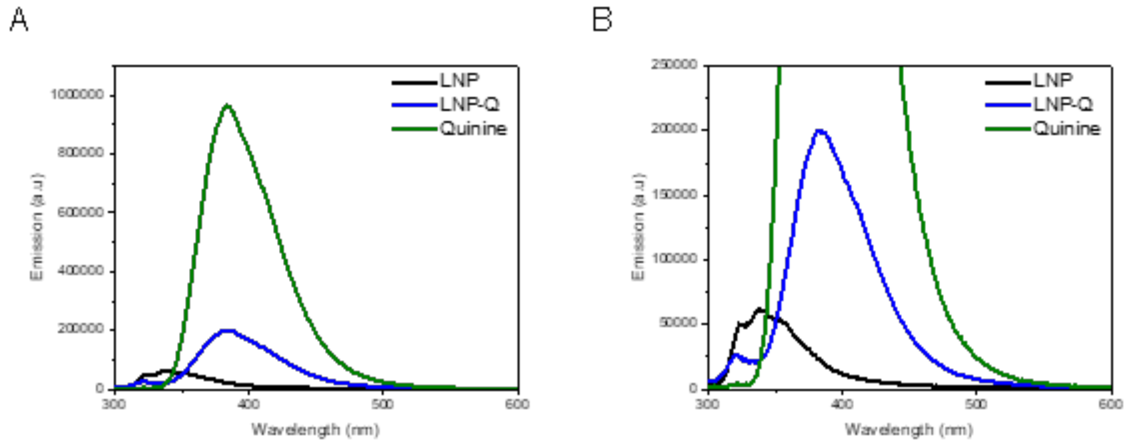


***Figure S24:*** *(A) Fluorescence emission spectra of drug-free LNP and LNP loaded with quinine recorded at an excitation wavelength of 288 nm show an overlap between quinine and quinine-loaded LNP.* ***(B)*** *Enlarged fluorescence spectra. Measurements were conducted at room temperature using a slit width of 3 nm.*
